# Functional Relations of CSLD2, CSLD3, and CSLD5 Proteins during Cell Wall Synthesis in *Arabidopsis*

**DOI:** 10.1101/2023.04.25.538313

**Authors:** Jiyuan Yang, Abira Sahu, Alexander Adams, Shuyao Yin, Fangwei Gu, Heather B. Mayes, Erik Nielsen

## Abstract

Plant cell expansion is a dynamic process that is physically constrained by the deposition of new cell wall components. During tip growth, new cell wall materials are delivered at a restricted plasma membrane domain which results in a highly polarized expansion within that specified domain. Previous studies demonstrated that this process requires the activities of members of the *Cellulose Synthase-Like D (CSLD)* subfamily of cell wall synthases. CSLD3 displays β-1,4 glucan synthase activity, but whether other members of CSLD subfamily share this conserved biochemical activity, and whether CSLD proteins form higher-order complexes to perform β-1,4 glucan synthase activities have not been determined. Here, we use genetic methods to demonstrate that CSLD2 and CSLD3 functions are interchangeable during root hair elongation and cell plate formation, while CSLD5 provides a unique and irreplaceable function in the formation of cell plates. Importantly, genetic analysis with inactivated versions of CSLD3 show that, unlike CESA proteins, CSLDs do not require the simultaneous presence of different isoforms to perform catalytic cell wall synthase activities. *In vitro* biochemical activity experiments confirmed that CSLD2, CSLD3, and CSLD5 proteins displayed β-1,4 glucan synthases activities. Taken together, these results indicated that while all three vegetatively expressed CSLD proteins possess conserved β-1,4 glucan synthase activities, that CSLD5 has a more complicated and specialized role during cell plate formation.

## INTRODUCTION

Plant cells are embedded within a load-bearing extracellular matrix, the cell wall. To change the size and shape of plant cells during growth and development, new cell wall material must be properly delivered and incorporated into existing cell walls during cell expansion. Two major mechanisms that control changes in plant cell shape are called diffuse growth and tip growth (Mathur and Hülskamp, 2001). During diffuse growth, the primary load-bearing components of cell wall, cellulose microfibrils, are synthesized directly at the plasma membranes by large integral membrane protein complexes called cellulose synthase complexes (CSCs), comprised of multiple catalytic subunits encoded by CESA (Cellulose Synthase) proteins (Cosgrove, 2005; Lerouxel et al., 2006). The new cellulose microfibrils are deposited and integrated in ordered arrays along the entire expanding faces of the cells, which are often transversely oriented to the major axis of cell expansion (Grierson et al., 2014). On the other hand, during tip growth, new cell wall material is selectively delivered and deposited by polarized secretion only at restricted plasma membrane domains within the cell, resulting in a highly polarized cellular expansion (Hepler et al., 2001; Cole and Fowler, 2006).

Root hairs are tip-growing tubular extensions of root epidermal cells, which play an important role in the uptake of water and nutrients (Peterson and Farquhar, 1996; Jungk, 2001). Root hairs are formed by two separate processes: bulge initiation and subsequent tip growth (Vissenberg et al., 2001). In *Arabidopsis*, the initiation of bulge formation in root hair cells typically occurs at the apical end (closest to the root apical meristem) of the root hair precursor cell (Schiefelbein and Somerville, 1990). This initial bulge formation is thought to be regulated by the establishment of an apically focused auxin gradient within the root hair cell precursor (Molendijk et al., 2001; Jones et al., 2002; Singh et al., 2008), and appears to be primarily regulated by cell wall expansion rather than incorporation of new cell wall materials (Jones et al., 2002; Sampedro and Cosgrove, 2005; Lin et al., 2011). Once the bulge is formed, new polarized cell wall deposition is initiated and confined to the expanding tip of the growing hair, leading to the elongated outgrowth (Schnepf, 1986; Foreman and Dolan, 2001; Singh et al., 2008; Shibata and Sugimoto, 2019). During this process, Golgi-derived secretory vesicles containing polysaccharides such as hemicellulose, pectin, and cell wall proteins are enriched in the subapical cytoplasm in the growing hairs, called the “vesicle rich zone” (Grierson et al., 2014). These cell wall cargoes are released to the extracellular matrix through exocytosis and are incorporated into the newly forming cell wall. Cellulose is also deposited within this apical region in tip-growing root hair cells and cellulose synthase inhibitor treatment results in the termination of elongation and root hair rupture (Park et al., 2011), indicating cellulose synthesis and deposition at the growing tip is essential for root hair elongation. Electron microscopy confirmed that short, randomly arranged cellulose microfibrils could be observed in the initial cell wall layer at the tip region (Newcomb and Bonnett, 1965; Galway et al., 1997; Emons and Mulder, 2000). An additional inner cell wall layer was found ∼25 um from the growing tip (subapical region), which contains parallel arrays of cellulose microfibrils that organized in a helical orientation along the length of root hair (Emons and Wolters-Arts, 1983; Emons, 1994; Emons and Mulder, 2000). In the mutants of *cesa1*, *cesa3*, and *cesa6*, which are required for the cellulose synthesis of the primary cell wall, tip-growing root hairs are formed and no significant defects in tip-restricted elongation were observed (Park et al., 2011; Gu and Nielsen, 2013). Additionally, both EYFP-CESA3 and EGFP-CESA6 proteins are excluded from the apical plasma membranes of growing hairs, and no accumulation of these proteins is observed within the “vesicle rich zone” population of Golgi-derived secretory vesicles (Park et al., 2011). Taken together, these results suggest that CESA protein activities are not required for cellulose synthesis within the apical plasma membrane domains of tip-growing root hair cells.

Interestingly, in *csld3^kjk-2^* mutant plants root hair elongation is abolished, with root hair precursor cells undergoing cell rupture upon transition to tip-restricted elongation, resulting in a hairless phenotype in *csld3 ^kjk-2^* mutants (Favery et al., 2001). Furthermore, functional eYFP-CSLD3 fusion proteins are enriched in tip-localized apical plasma membranes, and specifically accumulate in transport vesicles that accumulate in the “vesicle rich zone” of actively growing root hair cells (Park et al., 2011). Furthermore, complementation of a *cesa6* mutant with a chimeric CESA6 protein containing a CSLD3 catalytic domain showed that these chimeric proteins assembled into CSC complexes with other CESA proteins, rescued the growth and cellulose defects of the *cesa6* mutant, and proteoliposomes containing purified, detergent-solubilized CSLD3 proteins could specifically utilize UDP-glucose as enzymatic substrates and synthesize endo-β-1,4-glucanase sensitive polysaccharides (Yang et al., 2020). Taken together, these results supported that the biochemical activity of the CSLD3 represents a UDP-glucose-dependent β-1,4-glucan synthase.

Both *CESAs* and *CSLDs* belong to the *Cellulose Synthase-Like (CSL)* superfamily, whose members are predicted to encode processive glycosyltransferases that synthesize plant cell wall polysaccharides (Fry et al., 2008). In *Arabidopsis*, there are seven CSL families including CSLA, CSLB, CSLC, CSLD, CSLE, CSLG, and CESA, among which the CSLD family shares the highest protein sequence similarity to CESA family (Richmond and Somerville, 2000). Functions of several CSL proteins have been determined in recent years. CSLA proteins are shown to catalyze the β-1,4-linkage between mannose subunits and are required for the synthesis of the backbone of mannan and glucomannan hemicelluloses (Liepman et al., 2005; Goubet et al., 2009). AtCSLC4 contributes to the synthesis of the β-1,4-glucan backbone of xyloglucan in the Golgi apparatus in *Arabidopsis* (Cocuron et al., 2007). However, differential localization of distinct CSLC proteins has been observed in barley (*Hordeum vulgare*). While *HvCSLC3* was coordinately expressed with putative xylosyltransferase genes, consistent with *AtCSLCs* function in biosynthesis of xylogucan, HvCSLC2 proteins localized to plasma membranes, and were not observed in the Golgi apparatus as would be expected if they contributed to xyloglucan biosynthesis (Dwivany et al., 2009). This observation raised the possibility that different CSLC family members may provide distinct biosynthetic activities, and that not all members of a particular CSL subclade may synthesize the same types of polysaccharides.

In *Arabidopsis*, there are six members in the CSLD subclade. *CSLD6* is currently thought to be a pseudogene and has not been found to be expressed in any plant tissues to date (Bernal et al., 2008). *CSLD1* and *CSLD4* are specifically expressed in pollen, and *csld1* and *csld4* mutants are male sterile due to defects in tip-restricted pollen tube elongation (Wang et al., 2001). The remaining *CSLD2, CSLD3,* and *CSLD5* genes are broadly expressed in vegetative tissues (Brady et al., 2007). Similar to *csld3 ^kjk-2^*, *csld2* mutants also display shorter and irregular root hairs, indicating CSLD2 is also important for proper tip-restricted root hair growth (Bernal et al., 2008). The functional roles of CSLD cell wall synthases are not restricted solely to cells undergoing tip-restricted expansion. While *csld5* mutants display no root hair defects, they have measurably shorter roots, smaller rosettes, and slightly reduced overall stature when compared to wild type (Col-0) (Bernal et al., 2007), additionally CSLD5, along with CSLD2 and CSLD3, all participated in the construction of new cell plates during plant cytokinesis (Gu et al., 2016). Cellulose deposition has been observed in newly forming cell plates (Miart et al., 2014). However, the predominant cell wall polysaccharide that accumulates at cell plates is likely the β-1,3-glucan, callose (Meikle et al., 1991; Samuels et al., 1995; Ferguson et al., 1998; Chen and Kim, 2009; Drakakaki, 2015). Given that members of some CSL clades, notably CSLCs which accumulate in distinct compartments (Cocuron et al., 2007; Dwivany et al., 2009) may have distinct biochemical activities, we decided to test to what degree different CSLD proteins were functionally interchangeable. Additionally, at least three distinct CESA proteins are needed in order to assemble functional CSCs during synthesis of cellulose microfibrils (Arioli et al., 1998; Taylor et al., 1999; Fagard et al., 2000; Taylor et al., 2000; Scheible et al., 2001; Taylor et al., 2003; Desprez et al., 2007; Persson et al., 2007; Taylor, 2007). We therefore also wanted to determine whether, like CESA proteins, multiple CSLD isoforms were required to assemble functional cell wall synthase complexes *in vivo*, and directly tested the biochemical activities of multiple CSLD family proteins. To test these questions, we used genetic methods to demonstrate that CSLD2 and CSLD3 proteins are functionally interchangeable with one another during root hair elongation and cell plate formation. While ectopic expression of *CSLD5* in root hair cells could partially rescue *csld3 ^kjk-2^* root hair elongation defects, this rescue was directly linked to the relative accumulation of this protein which is rapidly degraded in non-dividing cells. Furthermore, while CSLD2 and CSLD3 gene products were largely interchangeable, CSLD5 displayed a unique function during cell plate formation that neither CSLD2 nor CSLD3 could functionally replace. Finally*, in vitro* biochemical activity experiments revealed that CSLD2, CSLD3, and CSLD5 proteins all displayed conserved β-1,4 glucan synthases activities.

## RESULTS

### Distinct *csld* mutants showed different degrees of vegetative growth defects

While the primary morphological defects of *csld2*, *csld3^kjk-2^*, and *csld5* mutants have been previously described as defective root hairs (*csld2* and *csld3^kjk-2^*), and overall reduced stature (*csld5*) (Bernal et al., 2007; Bernal et al., 2008; Yin et al., 2011; Yoo et al., 2012), a more careful examination of defects associated with these individual mutants has not been previously undertaken. We therefore performed an analysis of the growth characteristics of these three broadly expressed CSLD proteins in early stages of growth and development in single *csld* mutant backgrounds (Figure 1). To assess the overall size of plants, the length of rosette leaves in three-week old plants was assessed (Fig 1A and 1D). While at early stages of seedling development and vegetative growth *csld2* mutants showed indistinguishable overall sizes to wild type (Col-0), both *csld3^kjk-2^* and *csld5* mutants displayed a ∼15% reduction in the length of rosette leaves of three-week old plants (Figure 1D). Additionally, 5-day old seedlings of all three vegetatively expressed *csld* single mutants displayed a 10%∼20% reduction on root length (Figure 1B and 1E), which was consistent with the overall size reduction in the *csld3^kjk-2^* and *csld5* mutant seedlings. Among them, the *csld3^kjk-2^*mutants displayed the most severe root length reduction. To distinguish whether the reduction is due to early germination defects, or a result of sustained absence of CSLD proteins, we recorded the root lengths of these mutant lines over seven days after germination (Figure 1B, 1C, and 1E). Compared to wild type, all three single mutants display reduced root length at any time point, indicating that the growth reduction shown in *csld* mutants is due to an overall slower growth rate. These growth defects were enhanced in *csld2/3^kjk-2^*, csld2/5, or csld*3^kjk-2^*/5 double mutants (Figure 6A) resulting in a ∼60% reduction in root length of 5-day old seedlings, indicating that all three vegetative CSLD proteins played important and at least partially redundant roles in both leaf expansion and root elongation during early stages of seedling development. We also measured height, fresh weight, and seed weight of the *csld* single and double mutants in five-week-old plants. We found that most *csld* lines, except *csld2* and *csld2/3^kjk-2^*, were significantly shorter and had lower fresh weight (Figure 2A and 2B). Intriguingly, *csld2* and *csld2/3^kjk-2^*double mutant plants displayed significantly taller inflorescences (Figure 2A); however, we did not find any difference in seed weight in any of the mutant lines (Figure 2C).

**Figure 1.**
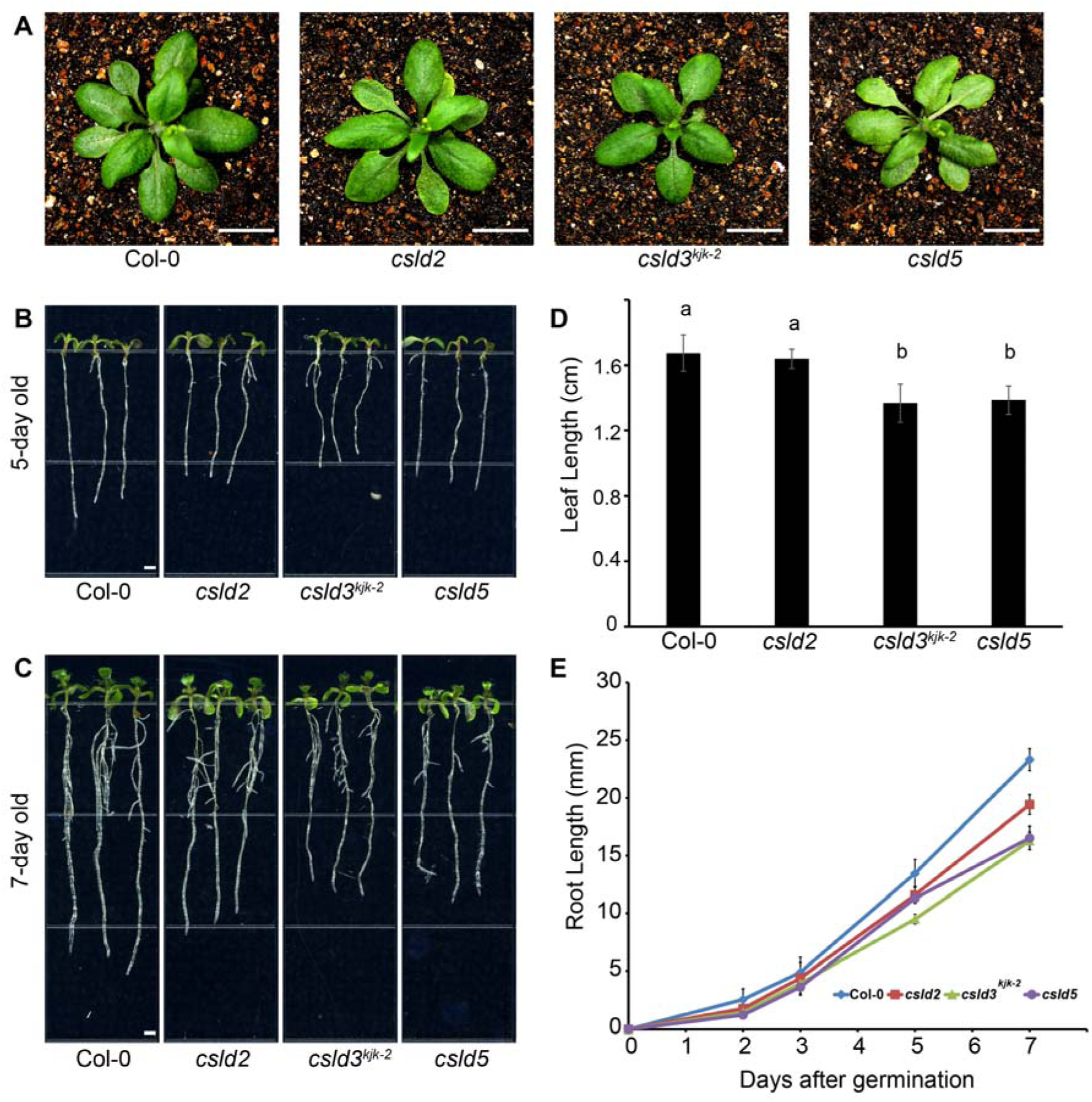
The single mutant of CSLDs displayed growth defects. **(A)** Rosette size of three-week old plants. *csld3^kjk-2^* and *csld5* mutants displayed smaller overall size of rosette leaves, while no significant change was observed in *csld2* mutants. The length of rosette leaves was measured and quantified in **(D)** (4-5 leaves per plant, 8 individuals). Five-day-old **(B)** and seven-day-old **(C)** seedlings were recorded using an Epson Perfection 4990 Photo scanner. The lengths of root regions were measured using Fiji-ImageJ (Schindelin et al., 2012). *csld2, csld3^kjk-2^* and *csld5* single mutants displayed varying degrees of growth reduction in root expansion. The length of roots is recorded and quantified in **(E)**. The *csld* mutants displayed a slower growth rate compared to wild type (Col-0) (n = 15). Statistically significant differences between group means were computed by ANOVA and Tukey’s HSD and they are denoted by lowercase letters. Groups that are significantly different from each other (p<0.05) are indicated by different sets of letters.

**Figure 2.**
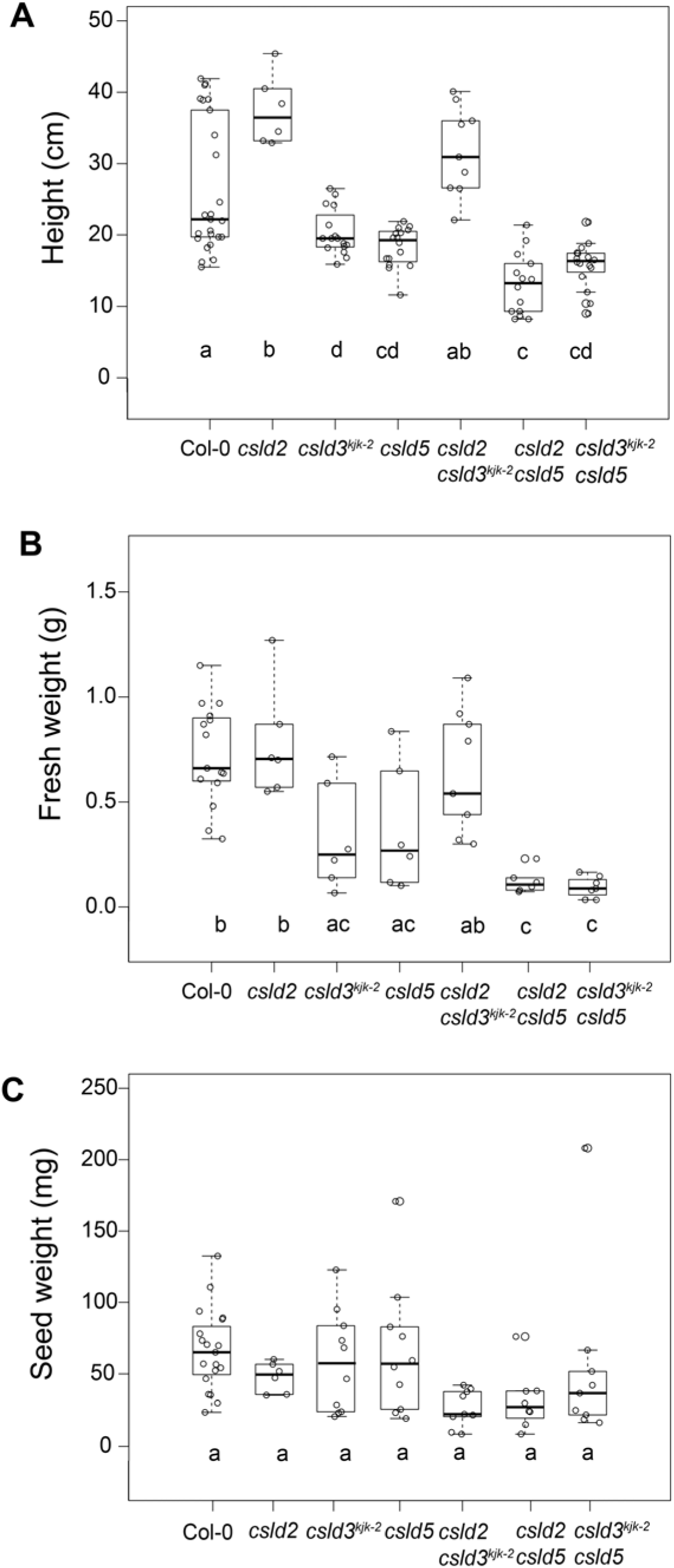
Phenotypic characterization of mature csld single and double mutants. Height (**A**), fresh weight (**B**), and seed weight of 5-week-old plants were measured. Although there was no significant difference in seed weight, most *csld* mutants showed reduced height and fresh weight compared to WT. Data were collected from 6-25 plants. Whiskers of the box plots represent 95% confidence interval. Statistically significant differences between group means were computed by ANOVA and Tukey’s HSD and they are denoted by lowercase letters. Groups that are significantly different from each other (p<0.05) are indicated by different sets of letters.

To investigate whether the reduced organ size observed in rosette leaves and roots was due to cell division or cell elongation, we quantified the cell division index and cell length in elongation zones in growing roots (Figure S1). The cell elongation zone was defined from the first root cortical cell that has longer cell length than cell width to the appearance of the first epidermal root hair. The distance from the QC to elongation zone defined the cell division zone and this region of the root was used to calculate the cell division index. Interestingly, while *csld2* and *csld3^kjk-2^* mutants showed indistinguishable cell division indexes to wild type (Col-0), *csld5* mutant had a ∼12% reduction in number of cell division events in this region of the root (Figure S1B). More severe reduction was observed in *csld2/3^kjk-2^*, *csld2*/5, or csld*3^kjk-2^*/5 double mutants, albeit as additive effects. Cortical cell length was used as a cell elongation index, where *csld3^kjk-2^* displayed a ∼30% reduction vs wild-type plants, with even shorter cell lengths observed in *csld2/3^kjk-2^*, *csld2*/5, or csld*3^kjk-2^*/5 double mutants (Figure S1C). Taken together, these data indicated that in addition to defects in root hair formation, that *csld2*, *csld3*, and *csld5* mutants displayed various additional defects in cell division and cell elongation, particularly during formation of root tissues in *Arabidopsis*.

Previous results indicated that CSLD3 localized to apical plasma membranes in actively growing root hairs, while CESA proteins were excluded from this plasma membrane domain (Park et al., 2011). To examine this in more detail, we generated transgenic plants that expressed both fluorescently-tagged CSLD3 and CESA6 under their natural promoter sequences (Figure 3). We observed that EYFP-CESA6 was solely present in the cell body plasma membranes during the early stages of hair formation, while Cer-CSLD3 was exclusively found in the plasma membrane of newly forming hairs (Figure 3A, left panel). At later stages during root hair elongation, Cer-CSLD3 remained primarily in the apical PM domain, but we observed EYFP-CESA6 labeling in more distal plasma membrane domains in these tip-growing root hairs (Figure 3A, middle and right panels). This more distal EYFP-CESA6 localization, was consistent with where additional inner cell wall layers containing parallel arrays of cellulose microfibrils have been observed using TEM methods (Emons and Wolters-Arts, 1983; Emons, 1994; Emons and Mulder, 2000). When examining *csld* mutant root hair phenotypes, the root hair morphology in *csld5* mutants was indistinguishable to that of wild type (Col-0) (Figure 3B and 3C). This was consistent with the relatively low expression level of *CSLD5* in root hair cells (Brady et al., 2007), as well as reflective of the low levels of CSLD5 protein accumulation we would expect to observe in these non-dividing cells (Gu et al., 2016). Both *csld2* and *csld3^kjk-2^* mutant phenotypes were most prominent during tip-restricted root hair elongation as previously described (Favery et al., 2001; Wang et al., 2001; Bernal et al., 2008; Galway et al., 2011; Park et al., 2011; Hu et al., 2019). In *csld2* mutants, bulges were formed at the base of growing root hairs, and early termination of root hair elongation resulted in significantly shorter hair length compared to wild type (Col-0) (Figure 3B). In *csld3^kjk-2^* mutants, essentially no root hairs were observed (Figure 3B). These phenotypes were consistent with the expression patterns of *CSLD2, CSLD3* and *CSLD5* generated using the *Arabidopsis* eFP Browser (http://bar.utoronto.ca/efp/cgi-bin/efpWeb.cgi) (Figure S2 (Brady et al., 2007)). Both *CSLD2* and *CSLD3* displayed increased expression levels in cells in the elongation zone and in developing root hair cells. While *CSLD3* expression was elevated in both hair-forming and non-hair forming epidermal cells as well as cortical and endodermal cells in the elongation zone, *CSLD2* expression in this region was more highly restricted to hair-forming epidermal cells. Additionally, while only *CSLD3* expression was elevated during root hair initiation (region 8), *CSLD2* expression levels showed a second peak in expression in later stages of root hair cell elongation, while *CSLD3* did not. In contrast, consistent with its role in cell plate formation during cell division, *CSLD5* expression was elevated only in the meristematic zone.

**Figure 3.**
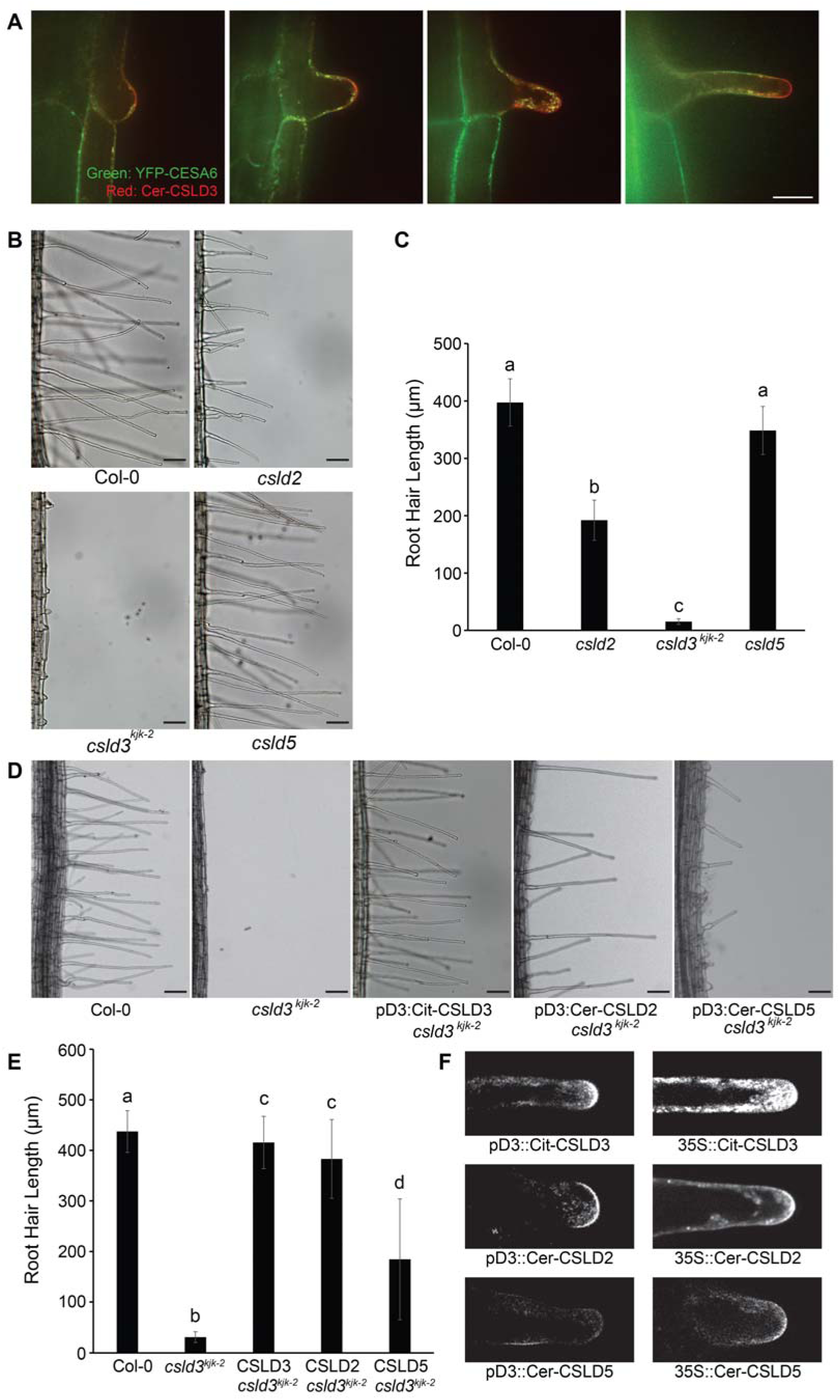
Ectopic expressed CSLD2 and CSLD5 driven by the *CSLD3* promoter promoted the root hair elongation. (**A**) The fluorescence of CESA6 and CSLD3 proteins was recorded in growing root hairs of five-day old seedlings. Green signals represent YFP-CESA6, Red signals represent Cer-CSLD3. The distinct localization from two colors represents the specific accumulation of CSLD3 at the apical of plasma membrane. (**B**) Root hair morphology analysis in three *csld* single mutants. Five-day-old seedlings were used to observe and record the phenotypes. In *csld2* mutants, the root hair length displayed a nearly ∼50% reduction compared to wild type (Col-0). Bulges were formed at the base of growing root hairs. In *csld3^kjk-2^*mutants, bulges were able to initiation, while nearly no root hair cells were able to continue the tip-restricted expansion, and results in a hairless defect. In *csld5* mutants, the morphological shape of root hair is not affected compared to wild type (Col-0). The length of root hairs (n=15) is quantified in **(C)**. **(D)** Root hair morphology analysis of stably transformed *csld3^kjk-2^* mutant plants expressing fluorescence tagged CSLD2, CSLD3, and CSLD5 under the endogenous CSLD3 promoter. When expressing Citrine-CSLD3, the hairless defects is fully restored, and root hair length is indistinguishable to wild type (Col-0). When expressing Cerulean-CSLD2, normal root hairs are generated as well. Shorter root hairs are observed in plant expressing Cerulean-CSLD5. The lengths of root hairs (n=15) in these plants are quantified in **(E)**. **(F)** Localization of fluorescence tagged CSLD2, CSLD3, and CSLD5 driven by the endogenous CSLD3 promoter (left three panels) and *35S* promoter (right three panels) in growing root hair cells. Statistically significant differences between group means were computed by ANOVA and Tukey’s HSD and they are denoted by lowercase letters. Groups that are significantly different from each other (p<0.05) are indicated by different sets of letters.

### CSLD2 and CSLD3 are functionally interchangeable in root hair cells undergoing tip growth

To examine whether the root hair tip-growth defects observed in the *csld2* and *csld3^kjk-2^* mutants were due to distinct genetic or biochemical functions between *CSLD2* and *CSLD3*, or simply the results of gene dosage effects due to differential expression patterns, we first tested whether *CSLD2* could functionally replace *CSLD3* during root hair elongation. We generated the stably transformed *csld3^kjk-2^*plants expressing a fluorescently tagged Cerulean-CSLD2 under control of the endogenous CSLD3 promoter. These transgenic plants quantitatively rescued the hairless defects in the *csld3^kjk-2^*plants (Figure 3D) and generated normal root hairs that were indistinguishable from wild type (Col-0) root hairs (Figure 3E), indicating the CSLD2 protein could functionally replace CSLD3 protein function during root hair elongation. Interestingly, in the stably transformed plant lines expressing Cerulean-CSLD5 driven by the endogenous CSLD3 promoter, we also observed limited levels of root hair formation (Figure 3D). However, root hair lengths were shorter, and the percentage of root epidermal cells in hair-forming positions that formed root hairs was dramatically lower with an average of about 40% compared to wild type levels of more than 95% (Col-0) (Figure 3E).

To determine if the fluorescently tagged CSLD proteins in these transgenic lines were accurately delivered to the tip region of the elongating root hair cells, we examined the localization of ectopically expressed CSLD2 and CSLD5 proteins by confocal microscopy (Figure 3F). Like EYFP-CSLD3, Cerulean-CSLD2 localized to the apical plasma membranes, as well as the compartments that accumulated in the vesicle-rich zone in the tips of growing root hairs, when expression was driven by CSLD3 promoter sequences. This tip-localized accumulation was observed in lines driving this expression either by the constitutive *35S* promoter or by endogenous CSLD3 promoter sequences (Figure 3F, middle panels). When driven by the endogenous CSLD3 promoter, the fluorescence signal of Cerulean-CSLD5 was barely detectable in root hair cells (Figure 3F, lower left panel), consistent with earlier observations that CSLD5 proteins are actively degraded in non-dividing cells in plants (Gu et al., 2016). When localization of fluorescent CSLD5 fusions was examined in lines where this fluorescent fusion was driven by the stronger, constitutive *35S* promoter, some accumulation of Cerulean-CSLD5 proteins was detected and displayed accumulation within the apical region of growing tip (Figure 3F lower right panel), a localization similar to those observed for fluorescent fusions of CSLD2 and CSLD3 proteins. These results would be consistent with CSLD2, CSLD3, and CSLD5 all having interchangeable functions during root hair tip growth may be interchangeable, but that the instability of CSLD5 in non-dividing root hair cells reduces the ability of this protein to rescue root hair tip-growth in the *csld3^kjk-2^* mutant background.

### Role of CSLD2, CSLD3, and CSLD5 in development of trichomes and pavement cells of cotyledons

We next explored the role of CSLDs in the development of other, non-tip growing cell types that still undergo high degrees of asymmetric expansion, such as trichomes and epidermal pavement cells of cotyledons. First, using scanning-electron microscopy, we examined trichome morphologies on the third and fourth primary leaves of 21d old Arabidopsis plants (Figure 4A-4G). Interestingly, both *csld2* and *csld3^kjk-2^* mutants displayed trichomes with reduced numbers of spikelets (Fig 4H). In both wild type (Col-0) and *csld5* mutants, more than 95% of trichomes had three spikelets (Figure 4H). However, in *csld2* mutants only 80% of trichomes had three spikelets, with 20% observed with only two spikelets, and in *csld3^kjk-2^* mutants, only 33% trichomes had three spikelets, with 58% having two spikelets, and 9% with only a primary trichome spike (Fig 4H). Since in tip-growing root hair cells, *CSLD2* and *CSLD3* appeared interchangeable, we wondered if there may also be functional redundancy between these two genes during trichome development. We found that *csld2/3^kjk-2^* double mutants also displayed 34% trichomes with three spikelets, 55% with two spikelets, and 10% with a single spike which was statistically indistinguishable to *csld3^kjk-2^*alone (Figure 4H). Similarly, in *csld2/5* double mutants, only 80% of trichomes had three spikelets, with the remaining 20% containing two or one spikelet, respectively (Fig 4H). Analysis of *csld3^kjk-2^/5* double mutants also resulted in trichome spikelet defect rates similar to *csld3 ^kjk-2^* alone (Figure 4H) while the trichome spikelet phenotypes of *csld2/5* mutants is similar to *csld2* mutants. The lack of any significant additive effect on trichome morphology defects in *csld2/3^kj-2k^* and *csld3^kjk-2^/5* double mutants, and the fact that *csld2/5* double mutant trichome defects were similar to *csld2* alone, suggested that while both *CSLD2* and *CSLD3* affect trichome development, *CSLD2* played a much less significant role than *CSLD3*.

**Figure 4.**
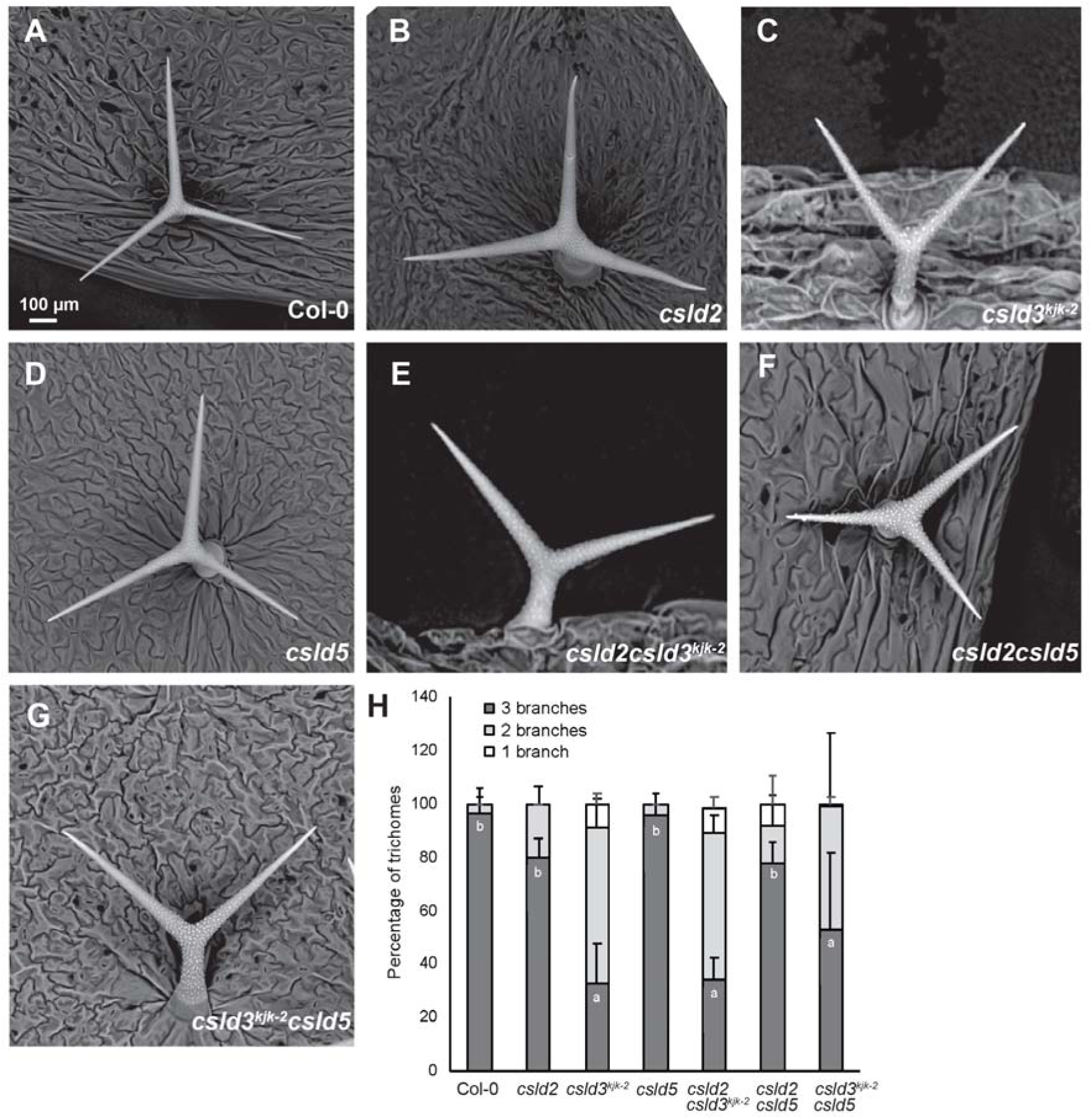
The presence of CSLD3 is important for trichome development in leaves. **(A-G)** Trichome morphology analysis in *csld* single and double mutants. Leaves from 3 week-old plants were used to observe and record the phenotypes using electron microscopy. Percent of trichomes with different degrees of branching are quantified in **(H)**. 8-12 leaves were imaged for each mutant and mean ± SD of each group is reported. Total number of trichomes analyzed range between 90-525 for each group. Statistically significant differences between group means were computed by ANOVA and Tukey’s HSD and they are denoted by lowercase letters. Groups that are significantly different from each other (p<0.05) are indicated by different sets of letters.

We next asked if there are morphological defects in the pavement cells of cotyledons in *csld* mutants. We used propidium iodide to stain the cell wall of cotyledons and imaged them using fluorescence microscopy. We then quantified lobe area normalized to total cell area and lobe number in the various *csld* mutants and double mutants (Figure 5). Number of lobes was determined using a modified version of the Lobefinder script (Wu et al., 2016; Thyssen et al., 2017), which we have called “LobePlotter” and it is available on https://github.com/xadams/LobePlotter (Figure S3).

**Figure 5.**
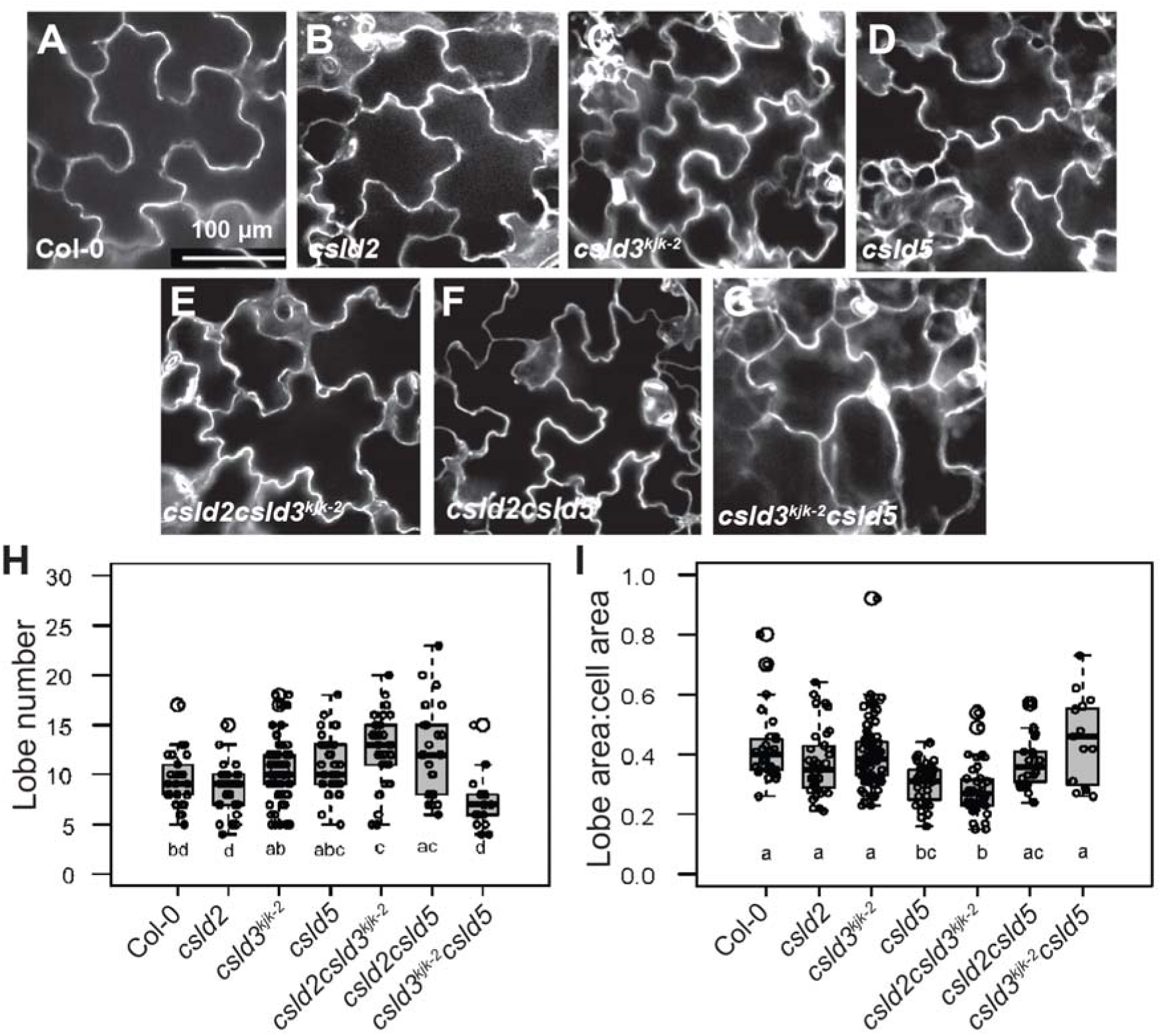
The development of pavement cells in cotyledons is affected in *csld* mutants. **(A-G)** The morphology analysis of leaf epidermal cells in *csld* single and double mutants. Cotyledons of 10-d old seedlings were used for imaging. Lobe number and area covered by the lobes are quantified in **(B)**. 17-37 cells were used for this analysis. Whiskers of the box plots represent 95% confidence interval. Statistically significant differences between group means were computed by ANOVA and Tukey’s HSD and they are denoted by lowercase letters. Groups that are significantly different from each other (p<0.05) are indicated by different sets of letters.

Our LobePlotter script functions the same as the original Lobefinder through Step 2 in which the program identifies points with locally maximized distance to a convex hull drawn around the epidermal cell. Lobefinder performs a refinement of the convex hull, whereas LobePlotter currently skips this refinement step. Lobefinder then derives the number of lobes and other global parameters from this refined convex hull. In contrast, the LobePlotter script iterates on these maxima to form a neck for each lobe based on connectivity from the neck point to points on the perimeter of the lobe. After determining the lobe necks, lobe-specific parameters (e.g. neck width, lobe height, and lobe width) can distinguished from the main body of the cell. These lobe-specific parameters can be calculated individually, and then averaged for each cell, or over multiple cells. Using this modified LobePlotter script we were able to directly measure the relative cell surface area contributed by cell lobes versus the central cell body. Although we did not detect any significant difference in lobe number between Col-0 and *csld* single mutants, number of lobes/cell was significantly higher in *csld2/3 ^kjk-2^*and *csld2/5* mutant plants (Figure 5H).

We also measured ratio of lobe area and cell body area to determine the cell area that is present in cell lobe structures (Figure 5I). The ratio of lobe area to total cell area was significantly lower in *csld5* and *csld2/3 ^kjk-2^* indicating that lobes represent a smaller fraction of the total cell surface area in these mutants. In the *csld2/3^kjk-2^* mutant, increased lobe number is associated with a lower ratio of cell area associated with these additional lobes, this quantification is consistent with large numbers of small lobes in this mutant background (compare panels 5A and 5E). Since this phenotype was observed in *csld2/3 ^kjk-2^* double mutant but not in the respective single mutants, *CSLD2* and *CSLD3* may play additive roles in lobe formation of pavement cells. Furthermore, *CSLD5* may also participate in the development of pavement cells as evident from the reduced lobe sizes observed in the *csld5* mutant. However, we did not find any significant difference in individual lobe area in *csld2/5* and *csld3 ^kjk-2^/5* double mutants. Further studies are required to dissect the roles of CSLDs and their mechanism of action during the development of pavement cells.

### CSLD5 plays a unique role during cell plate formation in plant cytokinesis

Earlier studies have shown that *csld5* displayed cell wall stub defects in root cortical cells and developing stomatal cells (Gu et al., 2016). These defects in *csld2/5* and *csld3 ^kjk-2^/5* double mutants were dramatically enhanced (Figure 6C and 6D), indicating that *CSLD2* and *CSLD3* are also involved in the formation of cell plates during plant cell division (Gu et al., 2016). Interestingly, although *csld2/3^kjk-2^* double mutants suffered enhanced growth reduction defects and displayed shorter roots similar to *csld2/5* and *csld3^kjk-2^/5* double mutants (Figure 6A and 6B), significant numbers of cell wall stubs were not observed in *csld2/3^kjk-2^* double mutants (Figure 6C). This result might indicate either that *CSLD5* performs a unique function during cell plate formation or, as with earlier results during root hair development (Figure 3), these *CSLD* genes may simply need to be expressed at the appropriate time during cell division with a *CSLD5* promoter sequence.

To investigate these possibilities, we generated stably transformed transgenic lines expressing Citrine-CSLD2 or Citrine-CSLD3 driven by *CSLD5* promoter sequences and examined if ectopically expressed fluorescent fusions of CSLD2 or CSLD3 could functionally replace CSLD5 and rescue *csld5* cell wall stub defects. We first transformed *csld5* mutants with Citrine-CSLD2 and Citrine-CSLD3 expressed behind the CSLD5 promoter, but these lines did not rescue *csld5* cell wall defects (Figure 7A upper panels). Statistically, in both cases, the percentage of cells displaying wall stub defects were indistinguishable from the *csld5* mutant alone (Figure 7B).

**Figure 6.**
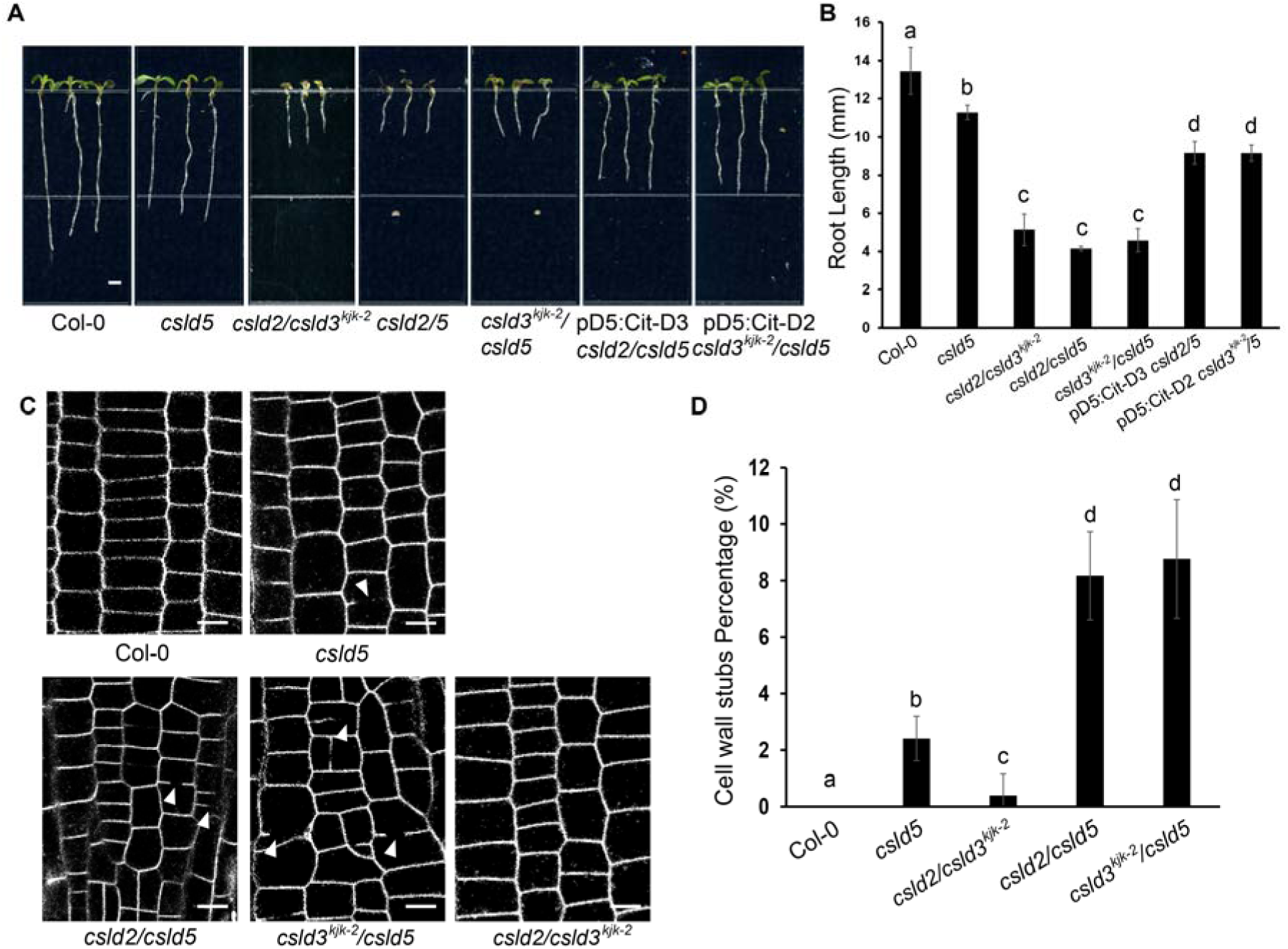
CSLD5 is essential and irreplaceable for cell wall deposition in cell plate formation during cytokinesis. **(A)** Root length analysis of five-day-old seedlings. The growth reduction of root tissue expansion is significant enhanced in *csld2/csld3, csld2/csld5,* and *csld3/csld5* double mutants. The length of roots was recorded and quantified in **(B)** (n=15)**. (C)** Confocal images of root epidermal cells from five-day-old seedlings. The seedlings were first stained with FM4-64 to visualize the edges of the root cortical cells. The cell division defects (cell wall stub) exhibited in *csld5* mutant are highly enhanced in *csld2/csld5, csld3/csld5* double mutants (white arrowheads). No cell wall stub is observed in *csld2/csld3* double mutants. The percentage of cell wall stubs is quantified in **(D)** (n=15). Statistically significant differences between group means were computed by ANOVA and Tukey’s HSD and they are denoted by lowercase letters. Groups that are significantly different from each other (p<0.05) are indicated by different sets of letters.

**Figure 7.**
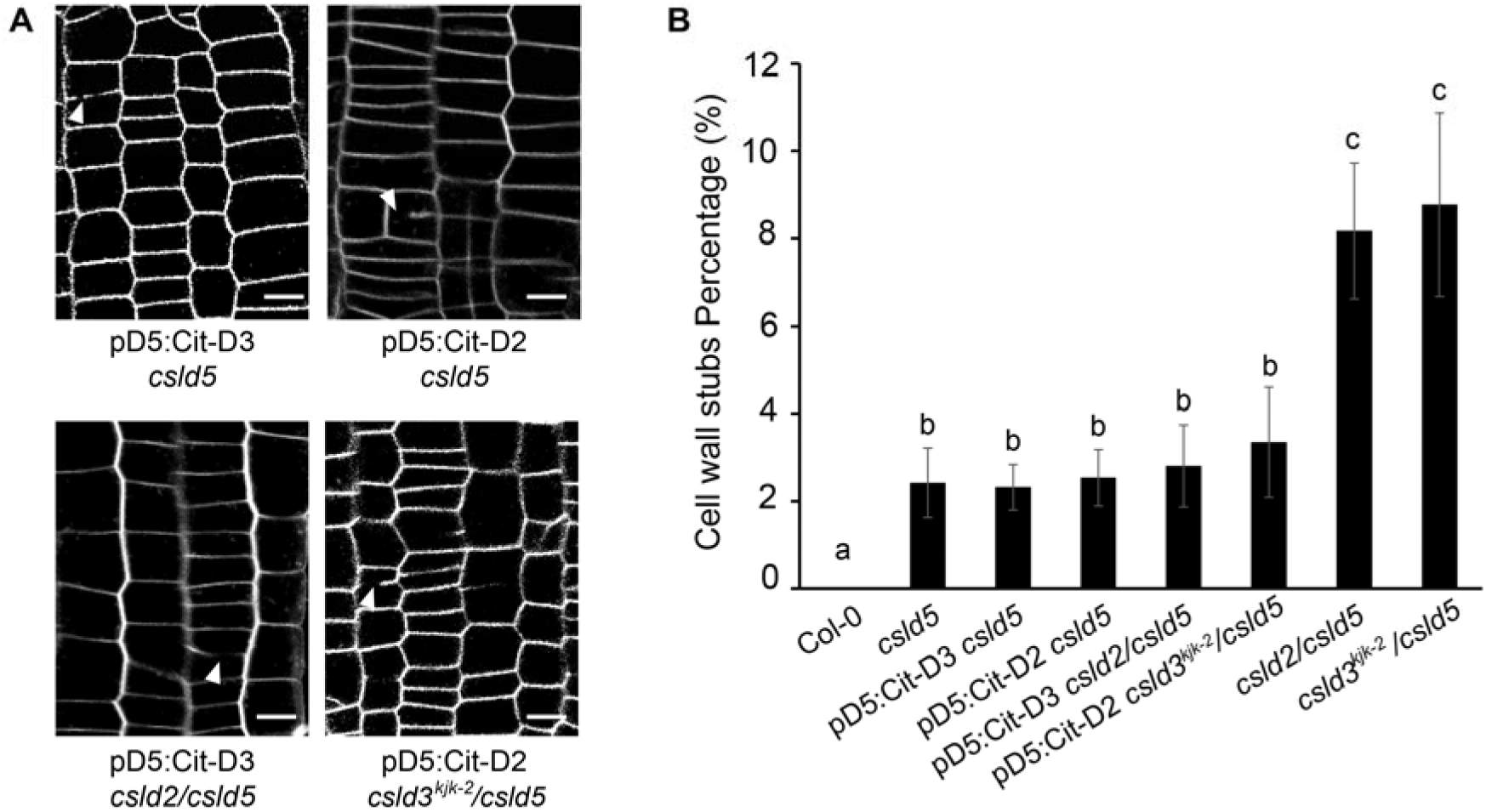
Neither Citrine-CSLD2 nor Citrine-CSLD3 can replace CSLD5 activity in *csld5* mutant backgrounds. **(A)** Confocal images of root epidermal cells from five-day-old seedlings. The cell division defects (cell wall stub) exhibited in *csld5* mutant cannot be restored in stably transformed plants expressing CSLD2 or CSLD3 driven by the endogenous CSLD5 promoter (upper two panels). When expressing CSLD2 or CSLD3 in *csld2/csld5, csld3/csld5* double mutant background, the presence of CSLD2 and CSLD3 could reduce the percentage of cell wall stubs compared to the parental *csld2/csld5, csld3/csld5* double mutants, and reaches to similar level to single *csld5* mutants. The percentage of cell wall stubs is quantified in **(B)** (n=15). Statistically significant differences between group means were computed by ANOVA and Tukey’s HSD and they are denoted by lowercase letters. Groups that are significantly different from each other (p<0.05) are indicated by different sets of letters.

Since *csld5* mutants displayed a relatively low percentage of cells in which cell wall stub defects were observed (Figure 6D), we also examined the propensity of Citrine-CSLD2 or Citrine-CSLD3 driven by *CSLD5* promoter sequences to rescue the more severe cell division defects observed in *csld2/5* and *csld3 ^kjk-2^/5* double mutant backgrounds (Figure 7A lower panels). When expressing Citrine-CSLD2 with the endogenous *CSLD5* promoter in *csld3 ^kjk-2^/5* double mutants, Citrine-CSLD2 should be the sole CSLD isoform present during cell plate formation. Quantification of the percentage of cell wall stubs present in these plant lines indicated that while ectopic expression of Citrine-CSLD2 could reduce the incidence of cell wall stubs versus the *csld3 ^kjk-2^/5* double mutant background, this reduction was only back to the levels observed in the *csld5* mutant alone (Figure 7B). Similarly, CSLD5 promoter-driven Citrine-CSLD5 in *csld2/5* double mutants again only reduced cell wall stub incidence back to levels observed in *csld5* mutants (Figure 7B). Taken together, these results indicate that ectopic expression of CSLD2 or CSLD3 fluorescent fusions behind a *CSLD5* promoter is insufficient to fully rescue the specific cell division defects observed in *csld5* mutants. Instead, CSLD2 and CSLD3 likely functionally contribute to cell plate formation in an interchangeable manner during cell plate formation, consistent with earlier observations of their roles in root hair tip-growth.

### Mutations in the TED motif alter or eliminate CSLD3 protein function *in vivo*

While the results of ectopic expression of CSLD2 and CSLD3 are consistent with these two proteins being functionally interchangeable (Figures 3, 6, and 7) they did not directly address whether CSLD proteins need to be catalytically active *in vivo* to rescue the corresponding mutant defects. To address this question, we took advantage of a recently published crystal structure of bacterial cellulose synthase (BcsA) to design catalytically dead versions of CESA and CSLD proteins (Omadjela et al., 2013; Slabaugh et al., 2014). Based on the BcsA structure, a trio of highly conserved amino acid residues, Threonine - Glutamic acid - Aspartic acid (TED) were identified. Within this TED motif, the aspartic acid has been proposed to function as a catalytic base during the formation of the β-1,4-glycosidic linkages during the synthesis of glucan polysaccharides (Morgan et al., 2013; Omadjela et al., 2013; Morgan et al., 2016). When examining the catalytic domains of the plant cellulose synthases, CESA and CSLDs, we found this TED motif is absolutely conserved in higher plant CESA and CSLD protein sequences (Figure S4).

To examine whether the glutamic acid and aspartic acid residues present in *Arabidopsis* CSLDs TED motifs might display some functional redundancy due to the similar chemical nature of their sidechains, we substituted the aspartic acid residue, and also the neighboring glutamic residue with alanine residues to remove the carboxylic groups of these potentially catalytic amino acids. These different mutant combinations of the TED catalytic motif were called “TEA” or “TAA” mutations, respectively. Stably transformed plants expressing Citrine-CSLD3-TAA with endogenous *CSLD3* promoter sequences (*pCSLD3*:Citrine-CSLD3-TAA) were unable to rescue the hairless defects in *csld3 ^kjk-2^* (Figure 8A and 8B). Interestingly, plants expressing Citrine-CSLD3-TEA driven by endogenous *CSLD3* promoter were able to generate short root hairs, and partially rescued the *csld3 ^kjk-2^* hairless defects (Figure 8B). Quantification of this partial rescue showed about 60% reduction of hair length in plants expressing the Citrine-CSLD3-TEA construct compared to wild type (Col-0), as well as the complementary rescue control (*pCSLD3*:Citrine-CSLD3) (Figure 8C). To assess whether either the early termination of root hair growth, or an overall slower growth rate, caused the shorter root hair phenotypes, we recorded the root hair growth in wild type (Col-0), *pCSLD3*:Citrine-CSLD3-TEA, and *pCSLD3*:Citrine-CSLD3-TAA expressing plants, and measured the length of growing root hair cells at different time points (Figure 8D and Supplemental Movie S1). Stably transformed plants expressing *pCSLD3*:Citrine-CSLD3-TEA showed a slower elongation rate compared to wild type (Col-0), while *pCSLD3*:Citrine-CSLD3-TAA expressing lines displayed essentially no root hair growth. From these results we concluded that a catalytically active form of CSLD3 was required to rescue loss of this protein in *csld3 ^kjk-2^*mutants, and additionally that unlike the bacterial cellulose synthase, BcsA, the glutamic acid in the TED motif appears capable of providing some residual level of activity in CSLD3.

**Figure 8.**
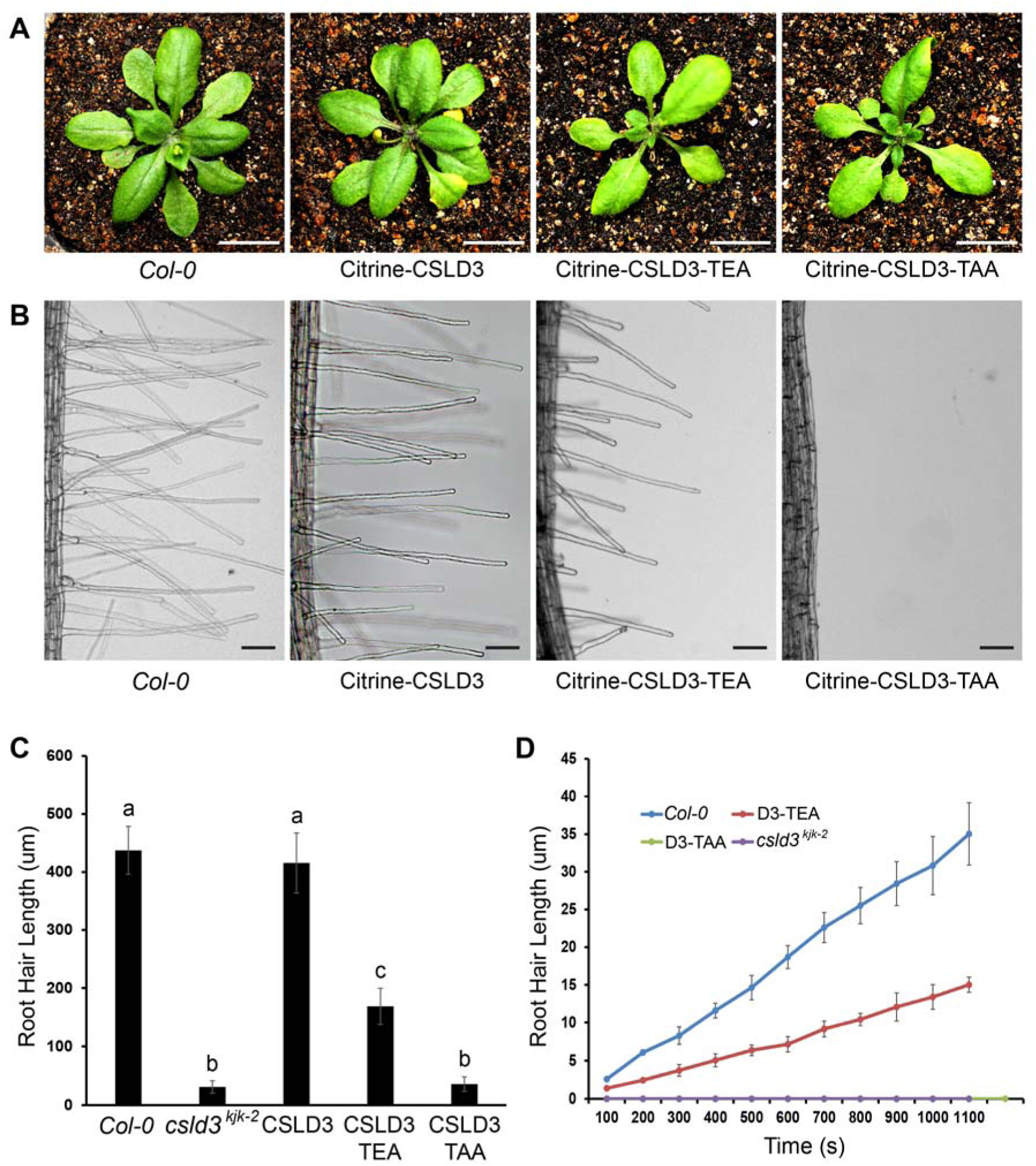
The quantitative rescue of *csld3^kjk-2^* hairless defects requires the catalytic activity of a β-1,4-glucan synthase. **(A)** Rosette size of three-week old plants. The stably transformed *csld3^kjk-2^* plants expressing Citrine-CSLD3 exhibits indistinguishable rosette size compared to wild type (Col-0). A smaller rosette size is observed in plants expressing Citrine-CSLD3-TEA and Citrine-CSLD3-TAA. The newly formed second layer of rosette leaves are still very small compared to wild type (Col-0), where the second layer of rosette leaves are in mature stage. **(B)** Root hair morphology analysis of five-day-old seedlings. The stably transformed *csld3^kjk-2^* plants expressing Citrine-CSLD3-TEA partially rescue the hairless defects and managed to generate root hairs are half long compared to wild type (Col-0). The length of root hairs is quantified in **(C)** (n=15). **(D)** The root hair length over time was recorded every 100 seconds for these plants (n=4). The slope represents the growth rate of root hairs. The stably transformed *csld3^kjk-2^* plants expressing Citrine-CSLD3-TEA display slower growth rate compared to wild type (Col-0), while plants expressing Citrine-CSLD3-TAA has no growth rate due to failures of promoting the tip-restricted expansion in root hair cells. Statistically significant differences between group means were computed by ANOVA and Tukey’s HSD and they are denoted by lowercase letters. Groups that are significantly different from each other (p<0.05) are indicated by different sets of letters.

To further distinguish if the partial rescue shown in plants expressing Citrine-CSLD3-TEA was due to reduced catalytic activity, or disruption of CSLD complex assembly directly, we examined the enzymatic activity of His-tagged CSLD3-TEA and CSLD3-TAA proteins *in vitro*. Using a yeast-based protein expression and purification system, proteoliposomes containing His-CSLD3-TEA proteins displayed a nearly four-fold increase in the K_m_ value for the UDP-glucose substrate (∼255uM), compared to wild-type His-CSLD3 which had a K_m_ value of ∼62uM for UDP-glucose (Figure 9A). The K_m_ value of CSLD3 for UDP-glucose was consistent when compared to previous CSLD3 and CESA6 K_m_ values (Yang et al., 2020). Consistent with the genetic rescue results, the His-CSLD3-TAA proteins failed to utilize the UDP-Glc as substrates and displayed only background levels of UDP formation when compared with blank (BLK) controls of proteoliposomes with no added CSLD proteins (Figure 9A). Taken together, these results are consistent with the conclusion that the quantitative rescue of *csld3 ^kjk-2^* hairless defects observed in seedlings expressing *pCSLD3*: Citrine-CSLD3 requires the catalytic activity of a β-1,4-glucan synthase, and that the partial rescue of root hair growth defects observed in lines expressing *pCSLD3*:Citrine-CSLD3-TEA is linked to reduced catalytic activity in these mutants.

**Figure 9.**
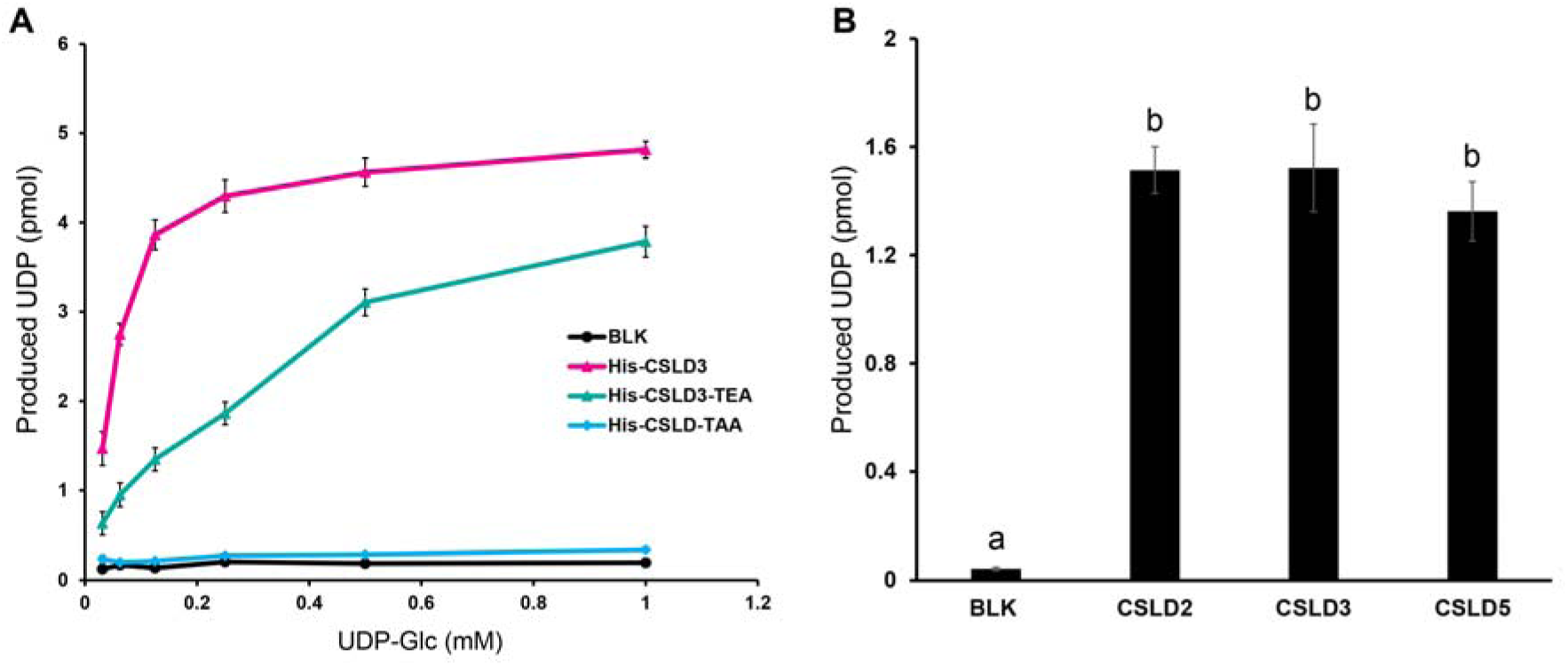
Purified CSLD2 and CSLD5 displayed β-1,4 glucan synthase activities *in vitro.* **(A)** Proteoliposomes containing His-CSLD3 (red line) and His-CSLD3-TEA (green line) displayed saturable UDP-forming activities when supplied with UDP-Glc. The Km values are 62 uM and 255 uM, respectively. Proteoliposomes containing His-CSLD3-TAA (yellow line) displayed no UDP-forming activities, similar to the negative control (BLK, black line) (n = 3). **(B)** Proteoliposomes containing His-CSLD2, His-CSLD3, and His-CSLD5 displayed similar UDP-forming activities when supplied with UDP-Glc (n=3). Statistically significant differences between group means were computed by ANOVA and Tukey’s HSD and they are denoted by lowercase letters. Groups that are significantly different from each other (p<0.05) are indicated by different sets of letters.

### *In vitro* activity experiments confirmed that CSLD2 and CSLD5 displayed β-1,4 glucan synthase activities

To further explore whether all vegetatively expressed CSLD proteins in *Arabidopsis* maintain the conserved β-1,4 glucan synthase activities, His-tagged CSLD2, CSLD3, and CSLD5 proteins were heterologously expressed in *S. cerevisiae*. To more specifically assess the enzymatic activities of these plant cell wall synthases, microsomal light membrane fractions were isolated from yeast expressing CSLD2, CSLD3, and CSLD5, and subsequently reconstituted into artificial proteoliposomes containing these purified cell wall polysaccharide synthases. These proteoliposomes were further incubated with UDP-Glc, to generate β-1,4 glucan polysaccharides. As previously demonstrated with CSLD3, both CSLD2 and CSLD5 also utilized UDP-Glc as substrates and generated free UDP with similar reaction rates to CSLD3 (Figure 9B), indicating all three vegetatively expressed CSLD proteins have conserved biochemical function as β-1,4 glucan synthases.

## DISCUSSION

Plant cells are surrounded by a rigid cell wall matrix. In order to grow and elongate, plant cells must synthesize new cell wall components and properly deliver these components and enzymes that modify or synthesize cell wall polysaccharides to specific sites where cell expansion is occurring (Richmond and Somerville, 2000; Cosgrove, 2005). In the past decades, the roles of CESA proteins in the synthesis of cellulose microfibrils, and the cellular mechanisms responsible for targeting these enzyme complexes to cortical microtubule arrays during diffuse growth has been well studied (Crowell et al., 2009; Gutierrez et al., 2009; Gu et al., 2010; Li et al., 2012; Liu et al., 2016; Zhu et al., 2018). On the other hand, the role that cellulose synthesis plays during construction of new cell walls and the mechanisms responsible for delivering enzymes involved in synthesis of this cell wall polysaccharide during highly focused cell wall expansion is still limited. Previous studies in our laboratory have shown that a member of the *CSLD* family, *CSLD3*, participates in the synthesis of a β-1,4-glucan polysaccharide during tip growth in root hair cells (Park et al., 2011; Yang et al., 2020). However, whether the other two vegetatively expressed CSLD proteins share a conserved biosynthetic activity, and how these three vegetatively expressed isoforms of the CSLD protein family cooperate during plant growth and development *in vivo* was not yet clear. To examine these questions, we first performed a detailed phenotypic analysis of growth and developmental defects of mutants of the three vegetatively expressed *CSLD* genes, *CSLD2*, *CSLD3*, and *CSLD5*. We found that *csld3 ^kjk-2^* and *csld5* mutants caused varying degrees of growth reduction in root elongation during early stages of seedling development (Figure 1), defects that were enhanced in *csld2/3^kjk-2^* and *csld2/5* double mutants (Figure 6). These early defects during seedling growth also resulted in altered stature and size in mature plants (Figure 2). Importantly, cell division and expansion were also broadly affected during development in these mutants (Supplemental Figure S1).

We also observed altered development of trichomes, epidermal pavement cells, and cortical root cells in *csld* single and double mutants (Figures 4 and 5). These observations indicated that CSLD proteins likely have additional roles in plant growth and development outside of their prominent roles in tip growing root hair cells and cell plate formation during cytokinesis. During the root hair tip-growth, stably transformed *csld3 ^kjk-2^* plant lines expressing *pCSLD3*: Cerulean-CSLD2 could fully rescue the *csld3 ^kjk-2^* hairless defects, indicating that *CSLD2* and *CSLD3* likely function interchangeably during cell wall deposition (Figure 3). Interestingly, partial rescue and shorter root hairs were observed in the stably transformed *csld3 ^kjk-2^* lines expressing *pCSLD3*: Cerulean-CSLD5 (Figure 3). We further compared the expression levels of fluorescently-tagged CSLD5 driven by different promoters (Figure 3F). Accumulation of Cerulean-CSLD5 was significantly enhanced in root hair cells when driven by the constitutive *35S* promoter compared to Cerulean-CSLD5 driven by endogenous *CSLD3* promoter sequences. These results suggested a gene dosage effect when rescuing the *csld3 ^kjk-2^* mutants: as long as a sufficient CSLD protein activity was present in the apical region of the growing tip, root hair elongation occurred normally. The partial rescue observed in the *pCSLD3*: Cerulean-CSLD5 expressing lines could be due to lack of accumulation of sufficient ectopically expressed CSLD5 in these non-dividing root hair cells. Unlike *CSLD2* and *CSLD3*, the expression level of *CSLD5* is highly regulated by the cell cycle (Gu et al., 2016). CSLD5 protein accumulation rapidly increases upon the initiation of M phase and is highly apparent during anaphase and telophase when cell plate formation occurs (Gu et al., 2016). Since CSLD5 protein level is regulated by APC^CCS52A2^ cell cycle regulatory complexes (Gu et al., 2016). The instability of CSLD5 protein and its degradation of CSLD5 in non-dividing cells might explain why ectopic expression of *CSLD5* could only partially restore root hair elongation, and significant accumulation of Cerulean-CSLD5 protein was only observed in partially rescued root hairs if expressed under control of the strong, constitutive *35S* promoter (Figure 3).

This interchangeable nature of *CSLD2* and *CSLD3*, but not *CSLD5*, was further supported by genetic rescue analysis of *csld2/5*, *csld3 ^kjk-2^*/*5* double mutant plants expressing *pCSLD5*: Citrine-CSLD3 and *pCSLD5*:Citrine-CSLD2, respectively. In *csld3 ^kjk-2^/5* double mutant plants expressing Citrine-CSLD2, CSLD2 is the sole CSLD protein isoform present in dividing cells. The ectopic expression of *CSLD2* causes a reduction in the cell wall stub percentage, but only to levels observed in *csld5* mutants. Similar results were obtained for *csld2/5* double mutant plants expressing Citrine-CSLD3. The fact that both plant lines displayed a reduction of cell wall stubs versus the parental double mutant backgrounds indicated that when a sufficient amount of CSLD protein is present, that both CSLD2 and CSLD3 are functionally interchangeable during cell plate formation. On the other hand, that no cell wall stubs were observed in *csld2/3 ^kjk-2^*double mutants might indicate that CSLD5 provides a unique function during plant cell division. Ectopic expression of either CSLD2 or CSLD3 under control of CSLD5 promoter sequences in *csld3^kjk-2^/csld5* and *csld2/csld5* double mutants only rescued the incidence of cell wall stubs back to levels observed in *csld5* mutant plants. These results indicated that even though CSLD2, CSLD3, and CSLD5 all participate in the process of cell plate formation, CSLD5 function is irreplaceable during cytokinesis. Based on these results, CSLD5, in addition to its beta-1,4-glucan synthase activity, might also serve as a “check point” protein during cytokinesis. The degradation and depletion of CSLD5 at the completion of cytokinesis may represent an M/S1 checkpoint for progression into new cycles of cell division. Taken together, the dosage-dependent rescue of CSLD proteins during root hair elongation and cell plate formation indicated that CSLD proteins, unlike CESA proteins, do not require the simultaneous presence of different protein isoforms to perform catalytic cell wall synthase activities (Gonneau et al., 2014; Hill Jr et al., 2014).

Previous results have shown that mutations of the “TED” catalytic core fully abolished the activity of CESA6 protein without affecting its ability to integrate into a higher-order complexes. CSCs containing the non-functional CESA6-TAA protein still displayed linear movement with a reduced speed (Yang et al., 2020). Due to the high degree of sequence similarity and overall protein topology between CSLDs and CESAs (Morgan et al., 2013; Slabaugh et al., 2014; Sethaphong et al., 2016), it remained unclear whether mutations in TED motif would abolish CSLD activity, and whether CSLD proteins could assemble into complexes containing other cellulose synthases. To test this, we generated stably transformed plant lines expressing the fluorescently tagged CSLD3-TEA and CSDL3-TAA proteins driven by the endogenous CSLD3 promoter. The TAA mutation fully abolished the biochemical activity of CSLD3 proteins *in vitro* (Figure 9), and plants expressing fluorescently tagged CSLD3-TAA proteins formed no root hairs (Figure 8). It is intriguing that plants expressing CSLD3-TEA proteins showed a rescue of approximately 50% of the root hair length. Further kinetics analysis demonstrated that CSLD3-TEA protein displayed higher K_m_ value, indicating reduced affinity for its substrate, UDP-Glucose. This result is consistent with the slower growth rate of root hairs observed in these plants (Figure 8). We also confirmed that both CSLD2 and CSLD5 display conserved β-1,4 glucan synthase activities that are the same as CSLD3 proteins (Figure 9).

Due to their significant sequence similarity, CSLD proteins were initially identified as pollen-specific “cellulose synthases” (Doblin et al., 2001). Recent demonstration that CSLD proteins, like CESA proteins, utilize UDP-glucose to synthesize beta-1,4-glucan polysaccharides, and that these proteins can assemble into higher-order complexes (Yang et al, 2020), raises the salient question of why plants may have evolved and maintained two distinct families of cell wall synthases that have similar biochemical properties. One possible explanation for this comes from the cell wall structures of root hairs, which contain two distinct cell wall layers. An initial cell wall layer accumulates at the rapidly growing tips (apical 20-30 μm) of these cells (Emons and Wolters-Arts, 1983; Emons, 1994; Emons and Mulder, 2000). Fibrillar cell wall elements, which are thought to represent cellulose microfibrils in this layer were typically shorter and displayed random arrangement (Newcomb and Bonnett Jr, 1965; Galway et al., 1997). Inner layer cellulose microfibrils, which appear ∼25 μm back from the root hair apex, were longer and often organized in a helical orientation along the length of the root hair, similar to what is observed in cells primarily undergoing diffuse expansion (Emons and Wolters-Arts, 1983; Emons, 1994; Emons and Mulder, 2000; Galway et al., 1997). It is generally accepted that cell wall expansion anisotropy is strikingly different at the apical regions of tip-growing cells (root hairs, pollen tubes) compared to diffuse growth in cell walls more distant from this apical expansion region of the cell (Baskin, 2005).

During diffuse expansion, new cellulose microfibrils are deposited in parallel arrays transverse to the main axis of cell expansion, and the high degree of anisotropy of these microfibrils ultimately leads to the asymmetric expansion in these cells (Dumais et al., 2004). The orientation of these cellulose microfibril arrays is controlled by the selective movement of cellulose synthase complexes along cortical microtubules arrays attached to the plasma membranes in these cells (Paredez et al., 2006). In contrast, in the tips of root hair cells cell wall expansion appears to be isotropic, which might be due to the random arrangement of the cellulose microfibrils at the tip of tip-growing cells (Newcomb and Bonnett Jr, 1965; Emons, 1994; Galway et al., 1997; Shaw et al., 2000; Dumais et al., 2004). Interestingly, cortical microtubule arrays are also absent in the tips of rapidly growing root hair cells, and treatment with microtubule depolymerization drugs did not induce root hair elongation defects (Van Bruaene et al., 2004). A soft, flexible and isotropically deposited cell wall is conducive to the development of root hair tip growth in the presence of rapid cell elongation. Similar to root hair elongation, in other specific plant growth processes like pollen tube extension and cell plate formation, the speed of cell wall deposition is the first priority compared to the rigidity of the overall structure. Instead of CESA cellulose synthases, CSLD cellulose synthases were observed in those development processes (Bernal et al., 2008; Park et al., 2011; Gu et al., 2016; Yang et al., 2020). The second layer of the inner cell wall, found in subapical regions of elongating root hairs might serve to strengthen the overall structural stability of these cell walls. Consistent with the presence of the second layer of the inner cell wall, cortical microtubules also appears in the subapical region (Van Bruaene et al., 2004), and CESA proteins localize to these subapical membranes in these regions of root hair cells. In cells in which rapid cell wall deposition or rapid expansion is required, the ability to deposit a soft and flexible cellulose microfibril network produced through the action of CSLD cell wall synthases might be favored. Alternatively, reinforcement, or modification, of existing cell walls might favor deposition of CESA-synthesized cellulose microfibril networks that display higher anisotropic organization or larger and more rigid cellulose mibrofibrils. The balance between “fast deposition” vs “rigid structure” might explain why plants appear to have maintained two closely related families of beta-1,4-glucan synthases (the CESA and CSLD protein families) that could meet different cell wall requirement in a variety of developmental processes associated with cell wall deposition and cell expansion.

## EXPERIMENTAL PROCEDURES

### Plant material and growth conditions

*Arabidopsis thaliana* lines used in this study were derived from *Col-0* ecotype. Seeds were sterilized with 10% Clorox bleach solution, rinsed five times with distilled water, then stored at 4°C for 2 days before being plated on growth medium comprised of 0.25X Murashige and Skoog Basal Medium,1% sucrose, and 0.6% phytagel. Plates were placed vertically in a growth chamber at 21°C and grown under long-day conditions (16 hours light (200 uE/m^2^s)/8 hours dark photoperiod). Three-day-old dark-grown seedlings were used for microscopy analysis. Five-day-old seedlings were used for morphology analysis. For propagation of mature plants, 14-day old seedlings were transferred to soil and grown in environmental chambers at 21°C under long-day conditions (16 hours light/8 hours dark photoperiod).

### Yeast expression plasmid construction and growth conditions

*S. cerevisiae* (Strain INVSc1, Thermo Fisher, Cat#: C81000) was used for protein expression. Untransformed yeast was cultured in YPD medium. Positive colonies containing pYES2/NT C plasmids (Thermo Fisher, Cat#: V825220), expressing N-terminal His-tagged CSLD2, CSLD3, CSLD5, CSLD3-TEA, were selected and cultured overnight at 30°C and 180 rpm in <u>SC-Ura + Glucose</u> medium composed of 1.9 g/L SC-Ura (uracil drop-out) powder, 1.7 g/L yeast nitrogen base without amino acids and ammonium sulfate, 5 g/L ammonium sulfate, and 20 g/L glucose. Yeast cells were harvested, rinsed in sterile water, and used to inoculate 200 ml of <u>SC-Ura + Raffinose</u> medium with the same nitrogenous base composition containing 20 g/L raffinose to an OD_600_ equal to 0.03. Cultures were grown for 14 to 16 h at 30°C and 180 rpm until the OD_600_ reached 2.0. Protein expression was induced by addition of 800 ml of <u>SC-Ura +</u> <u>Galactose</u> medium containing 20 g/L galactose, and cells were incubated for an additional 6 h at 30°C and 180 rpm. Yeast cells were harvested, weighed, flash frozen in liquid nitrogen, and stored at −80°C.

### Morphology analysis

The whole plants were pictured at three weeks old stage using Canon Kiss 5 digital camera. The raw images were collected at 5184*3456 pixel and cropped using Adobe Photoshop using the same scale bar level.

### Root hair and cell wall stubs morphology analysis

Root hair morphology characterization was carried out using a Nikon Eclipse E600 wide-field microscope with 4X (NA=0.2) and 10X (NA=0.3) Plan Apo lens as previously described (Kim et al., 2022). 15 individuals and over 100 root hairs were measured per line in total for quantification of root hair elongation.

Cell wall stub morphology analysis was acquired using a Leica confocal laser scanning microscope SP8 with a 40X Plan lens (NA=0.65). Seedlings were prepared as previously described (Gu et al., 2016). 15 individuals were measured per line.

### Root length, and Rosette size measurement

Images of 5-day-old seedlings were recorded using an Epson Perfection 4990 Photo scanner. The lengths of root regions were measured using Fiji-ImageJ (Schindelin et al., 2012). All transformed lines were grown side by side on the same plate, and at least 15 individuals were measured per line. Three independent biological replicates were performed for each line.

Images of three-week old plants were recorded using a Canon EOS R5 camera. The rosette size was measured using Fiji-ImageJ (Schindelin et al., 2012). All transformed lines were grown under same condition. For quantification, elder 4-5 rosette leaves were measured per individual, and at least 8 individuals were measured per line.

### Fluorescence microscopic imaging analysis

Images were acquired using a Leica confocal laser scanning microscope SP8 using a 60X oil lens (Type F Immersion oil, NA=1.518) and processed with the Leica Application Suite X (LAS X) Life Science Microscope software. YFP and citrine fluorescence were excited at 514 nm and visualized from 519 nm to 650 nm. Cell wall was stained using FM4-64, which was excited at 488 nm and visualized from 524 nm to 650 nm. Raw images were collected using 512*512 pixel images. Brightness and contrast were adjusted accordingly using Adobe Photoshop.

### Trichome imaging by scanning electron microscopy

Leaf 3 and leaf 4 of 21d old Arabidopsis seedlings were mounted on aluminum stubs (ϒ12.2 × 10 mm) with conductive carbon tape (8 mm wide) with their adaxial surface facing upwards. Leaf samples were coated with ∼15nm carbon using a carbon coater and then imaged under low pressure in JEOL JSM-7800 FLV Field Emission Scanning Electron Microscope. Images were captured at 140X magnification.

### Imaging of trichome branches using stereoscope

To quantify trichomes with defective branching, we imaged third and fourth leaves of 21d old Arabidopsis seedlings using a Zeiss stereo dissecting scope. Leaves were mounted on glass slide and glass brick was placed on the leaves to keep them flat. Images were captured at 2X magnification.

### Imaging pavement cells of cotyledons

Arabidopsis seedlings were grown in 0.25X Murashige and Skoog medium containing 1% sucrose and 0.6% phytagel for 10 days in a growth chamber (16 hours light (200 uE/m^2^s)/8 hours dark photoperiod). Before imaging, the seedlings were immersed in 10 ìg/ml propidium iodide solution then washed in distilled water. The seedlings were then placed on glass slide and cover slip was put on the cotyledons and taped tightly with the slide to keep the cotyledons flat. They were imaged using an Olympus spinning disk confocal imaging system equipped with an Andor iXon Ultra CCD camera and 20X oil immersion objective lens. Fluorescence from propidium iodide was detected using 488 nm laser excitation. To analyze the images, ROIs were drawn around individual cells using the polygon tool on ImageJ and lobe numbers were analyzed using a modified version of the Lobefinder script (Wu et al., 2016) available on https://github.com/xadams/LobePlotter. Lobe area: cell area was calculated using a Python script.

### Yeast protein extraction, purification, and proteoliposome reconstitution

5 g of yeast cells (corresponding to 8 L of <u>SC-Ura + Galactose</u> medium) expressing His-tagged CSLD3, CSLD3-TEA, CSLD3-TAA, CSLD2, CSLD5, or an empty vector were used for proteoliposome reconstitution. These experiments were performed as described previously (Yang et al., 2020).

### Accession Numbers

Gene information in this article can be found in The Arabidopsis Information Resource database (https://www.arabidopsis.org) under the following accession numbers: AtCSLD2 (AT5G16910), AtCSLD3 (AT3G03050), AtCSLD5 (AT1G02730), AtCESA6 (AT5G64740), and AtCSLA9 (AT5G03760).

## ACKNOWLEDGEMENTS

This article is based upon work supported by the National Science Foundation under grant no. 1817697 (E.N. and H.B.M).

---

**Figure S1.**
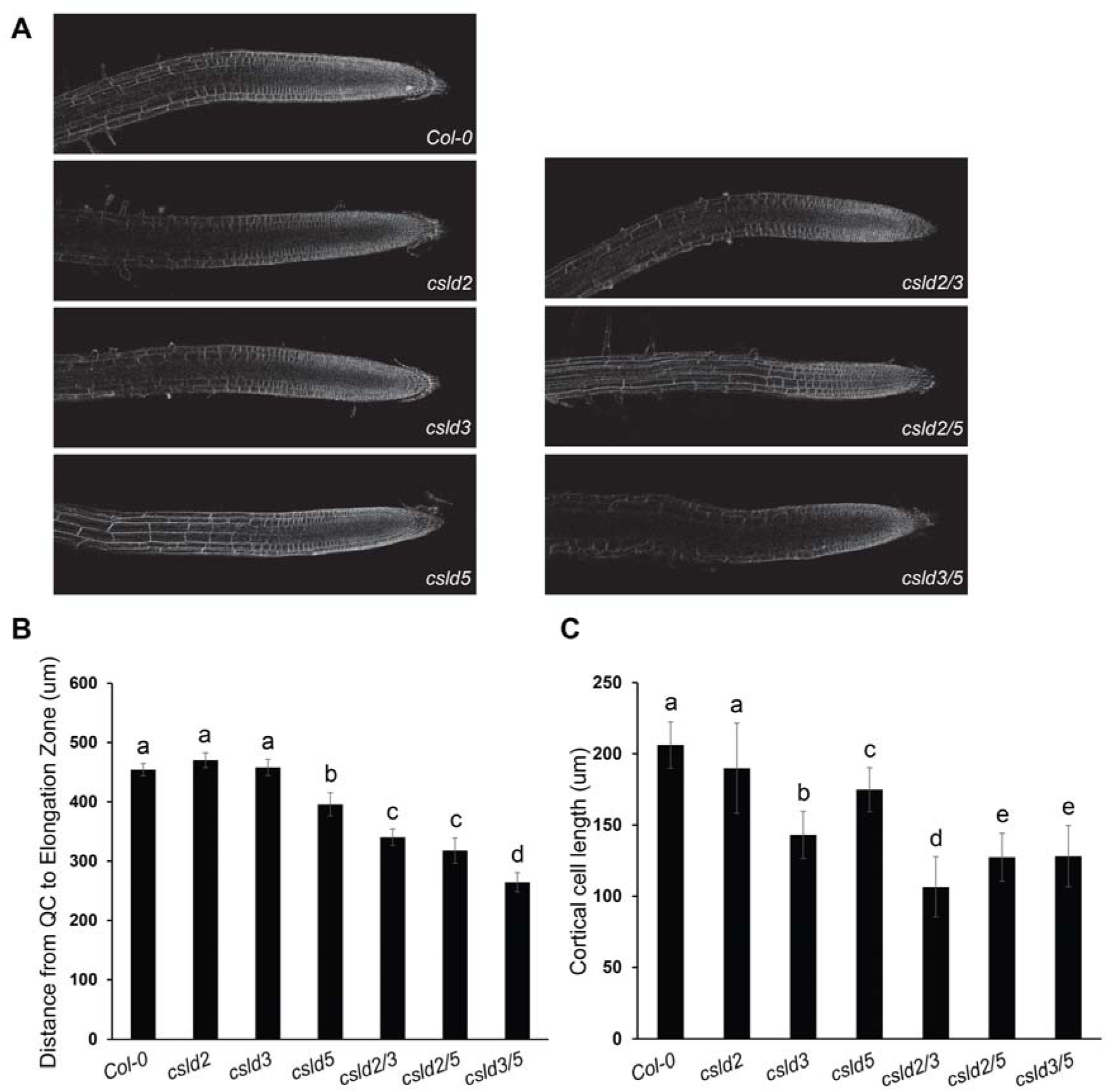
*csld* mutants displayed various effects on cell division and cell elongation in root tissue. **(A)** The morphology analysis of root cell length and cell number in *csld* single and double mutants. Five-day-old seedlings were used for imaging. Elongation zone is defined from the first cell that have longer cell length to cell width. Distance from QC to elongation zone define the cell division zone and was used as cell division index, quantified in **(B)**. Cortical cell length is quantified in **(C)** (n = 15). Statistically significant differences between group means were computed by ANOVA and Tukey’s HSD and they are denoted by lowercase letters. Groups that are significantly different from each other (p<0.05) are indicated by different sets of letters.

**Figure S2.**
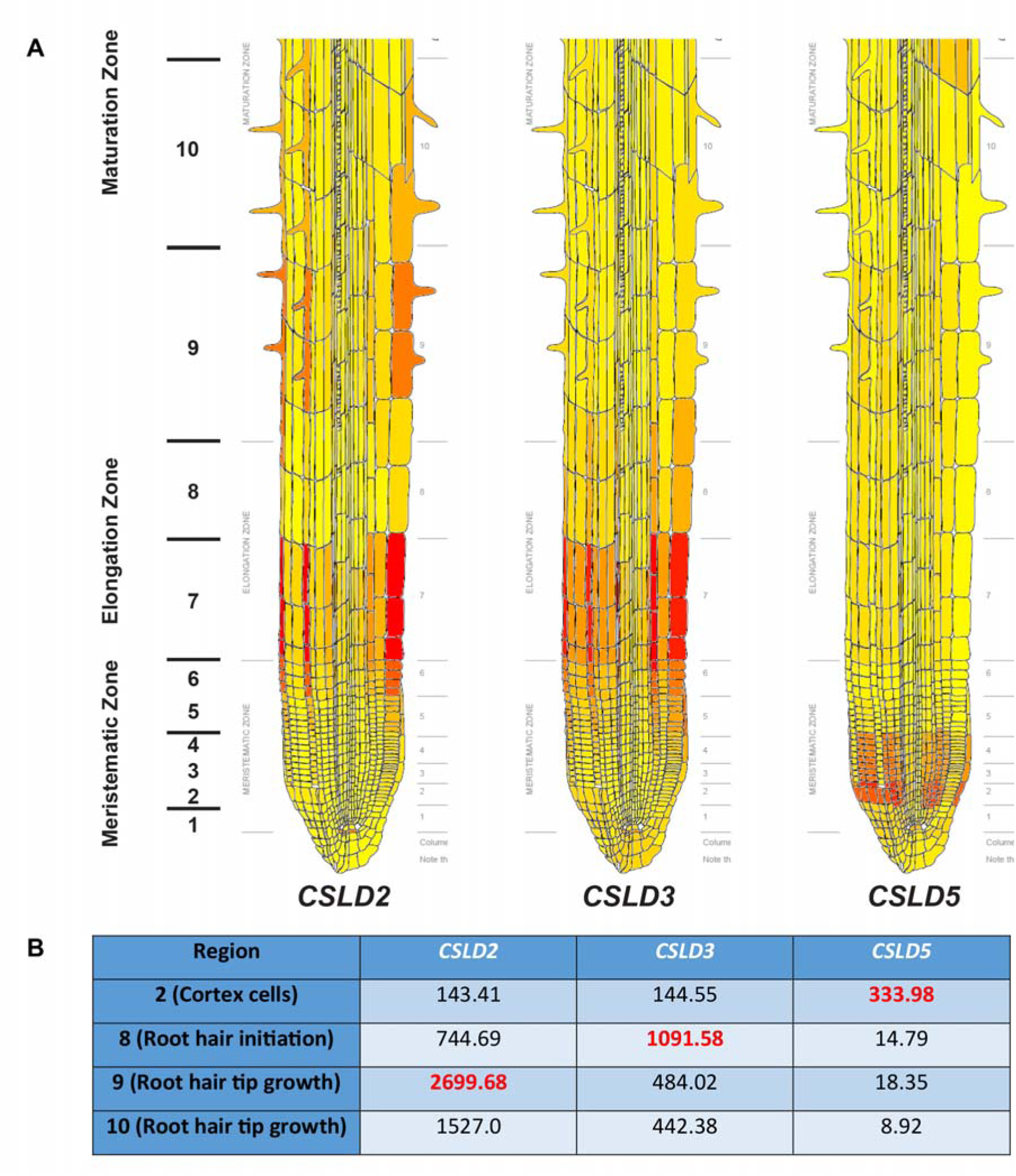
*CSLD2*, *CSLD3*, and *CSLD5* are differently expressed in root hair cells. **(A)** Root expression patterns of *CSLD2* (left), *CSLD3* (middle), and *CSLD5* (right) genes generated using the *Arabidopsis* eFP Browser (http://bar.utoronto.ca/efp/cgi-bin/efpWeb.cgi). Both *CSLD2* and *CSLD3* displayed increased expression levels in root hair cells, while CSLD5 was accumulated in root cortical cells meristematic zone (region 2) and displayed lowly expressed in root hair cells. *CSLD3* was expressed in hair cells initiating root hair growth (region 8), while *CSLD2* had high expression level after the root hairs were initiated (region 9). **(B)** Absolute expression level of *CSLD2*, *CSLD3*, and *CSLD5* in various regions. The highest values were highlighted in red.

**Figure S3.**
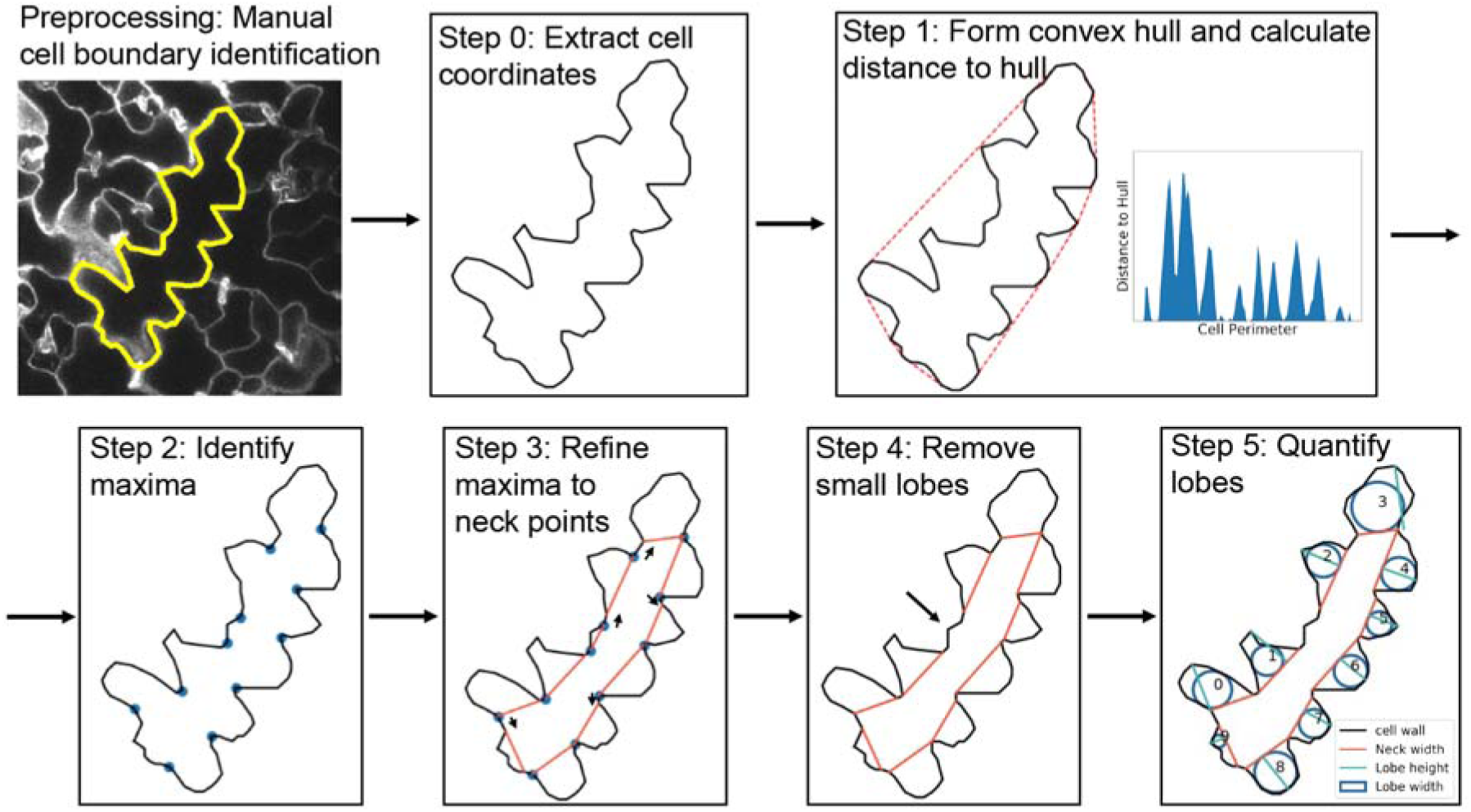
Schematic of the workflow of LobePlotter used to analyze the morphology of pavement cells of *csld* mutants. An ROI is drawn on the raw image to extract the cell coordinates. The following two steps are essentially the same as the Lobefinder which includes forming the convex hull then identifying points with locally maximized distance to the convex hull. Then our modified script iterates on these maxima to form a neck for each lobe, followed by calculating lobe specific parameters like neck width, lobe height, and lobe width.

**Figure S4:**
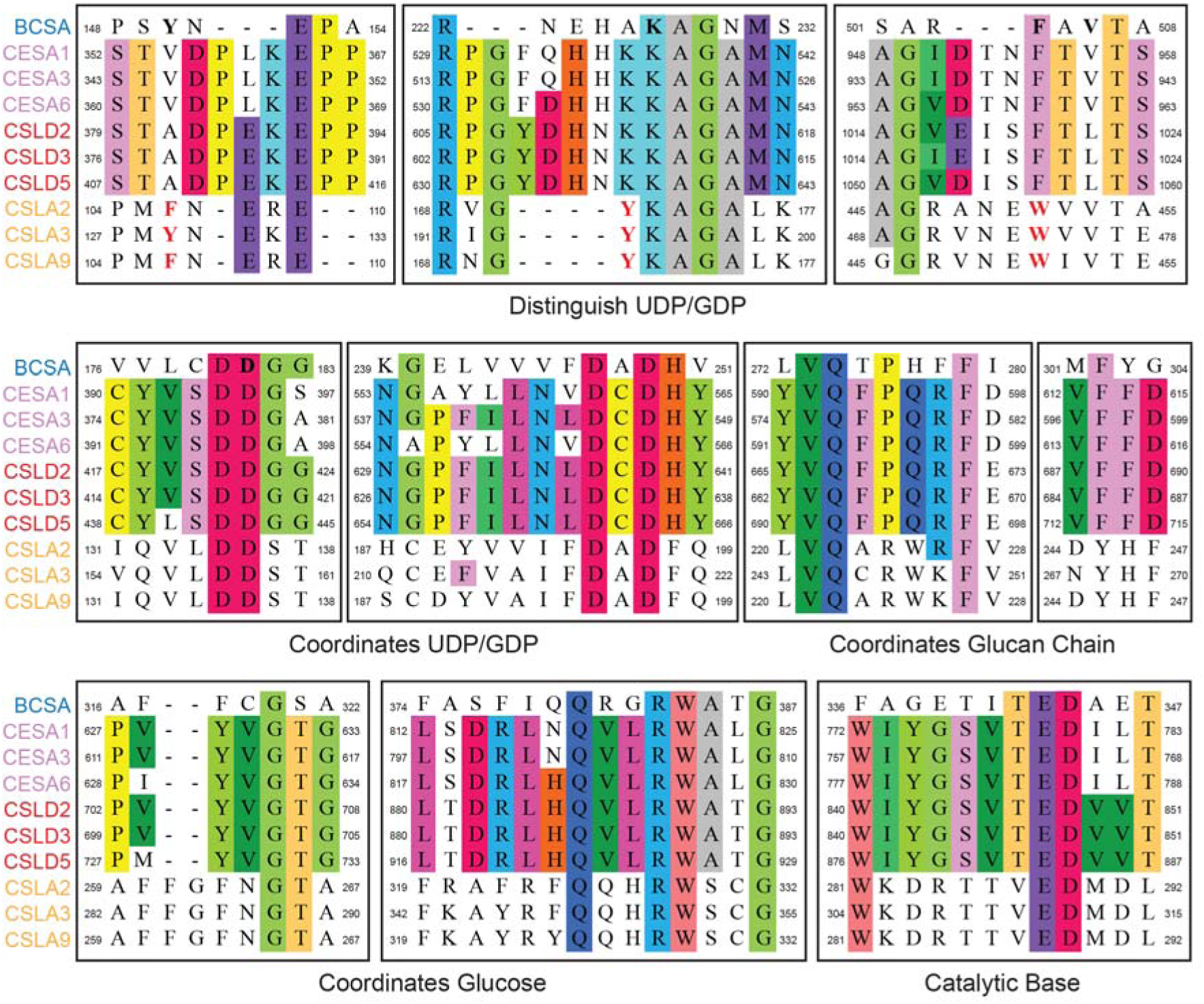
Alignment of amino acid residues of AtCESAs (pink), AtCSLDs (red) and AtCSLAs (yellow) was constructed by MEGA7 (Kumar et al., 2016). The amino acids in bold represent the identical sequences in three families including nucleotide coordinating regions (DD, DxD motifs), sugar coordinating regions (Q/RxxRW motif), and catalytic bases (TED motif). AtCSLD sequences showed higher identity to cellulose-synthesizing CESAs in these regions.

